# Stochastic Consolidation of Lifelong Memory

**DOI:** 10.1101/2021.08.24.457446

**Authors:** Nimrod Shaham, Jay Chandra, Gabriel Kreiman, Haim Sompolinsky

**Affiliations:** Center for Brain Science, Harvard University; Harvard Medical School; Center for Brain Science, Harvard University, Edmond and Lily Safra Center for Brain Sciences, the Hebrew University of Jerusalem

## Abstract

Humans have the remarkable ability to continually store new memories, while maintaining old memories for a lifetime. How the brain avoids catastrophic forgetting of memories due to interference between encoded memories is an open problem in computational neuroscience. Here we present a model for continual learning in a recurrent neural network combining Hebbian learning, synaptic decay and a novel memory consolidation mechanism. Memories undergo stochastic rehearsals with rates proportional to the memory’s basin of attraction, causing self-amplified consolidation, giving rise to memory lifetimes that extend much longer than synaptic decay time, and capacity proportional to a power of the number of neurons. Perturbations to the circuit model cause temporally-graded retrograde and anterograde deficits, mimicking observed memory impairments following neurological trauma.

## I. INTRODUCTION

Understanding the principles governing long-term memory is a major challenge in theoretical neuroscience. The brain is capable of storing information for the lifetime of the animal, while continually learning new information, so the brain must face the stability - plasticity dilemma: keep changing in order to learn new memories, but do so without erasing existing information. In humans, forgetting curves (retrieval probability vs. age of memory, sometimes referred to as retention curves), are found experimentally to be gracefully decaying with memory age, allowing for non-zero probability of retrieval for memories tens of years of age [1–4]. While retrieval probability curves monotonically decrease with memory age, retrievability of specific memories is non-monotonous with age, so that one might be able to retrieve a childhood memory, but forget events from last week.

Early attractor neural network models of long-term memory suffer from catastrophic forgetting: when the number of encoded memories is lower than a critical value, memories are retrievable with high precision, but when it is above that critical value, none of the memories can be retrieved [5–7]. Incorporating synaptic decay into the circuit enables continual learning, such that at any point in time recent memories are stable. However, the predicted forgetting curves exhibit a critical memory age, all memories newer than some age are almost perfectly retrievable, while all older ones are destroyed [8–13]. This is in contrast to the gracefully decaying forgetting curves in humans. Furthermore, the critical age is of the order of synaptic decay time, hence memories older than this time cannot be retrieved.

One of the main methods of studying the mechanisms of human memory is through memory disorders. Amnesic patients show a variety of patterns of forgetting. One is anterograde amnesia-reduced memory retrieval of events encoded after the onset of the disturbance to the circuit, presumably due to the inability to encode or store new memories. Another pattern is temporally-graded retrograde amnesia - when the probability of retrieval of memories encoded a short time before the pathology onset is lower than that of older events, giving rise to non-monotonic forgetting curves (an effect also known as Ribot’s law). Retrograde amnesia is typically explained by invoking memory consolidation theory, suggesting that memories must go through a stabilization process that is disrupted by the proximal onset of the disturbance [14–20]. In addition to possible cellular mechanisms, memory consolidation at the system level is mediated through a rehearsal process - reactivating memories in wakefulness or during sleep [21–25].

Several computational models have been proposed for memory consolidation through rehearsals [26–34]. However, all reported results were confined to a small number of memories; none demonstrated memory functionality and forgetting curves in a large circuit with a number of retrievable memories scaling with the number of neuron. None of the models obtain the scaling of capacity and memory lifetime with the number of neurons and other intrinsic parameters.

Here we present a neural network model for lifelong continual learning and memory consolidation. Our model continuously stores patterns of activity by Hebbian learning, and combines synaptic decay with stochastic nonlinear reactivation of memories. Our model generates intricate and rich memory forgetting behavior. Retrieval probability curves decay smoothly with memory age (exponentially or even as a power law), with characteristic times that can be orders of magnitude longer than the synaptic decay time. In addition, due to the stochasticity of the consolidation process, there is a large variability in the survival of individual memories of the same age. We show that at any given time, the number of retrievable memory scales linearly with the number of neurons, exhibiting adequate memory functionality expected for a robust neuronal circuit with distributed memories. Perturbations of the model circuit give rise to complex patterns of memory deficits,temporally-graded retrograde and anterograde amnesia, the details of which depend on the size as well as the nature of the perturbation.

Our theory relates global measures of memory functionality (memory capacity, characteristic memory lifetime) to intrinsic cellular and circuit parameters, such as synaptic decay rate and reactivation statistics, and provides new insight into how the brain builds and maintains the body of memories available for retrieval at each point in an animal’s life.

## II. RESULTS

### A. The model

Our model is based on the sparse version of the Hopfield attractor network model of associative memory [5, 6]. Memories are sparse ([7, 35–38]), uncorrelated *N* -dimensional binary activation patterns (*N* is the number of neurons) and are stored as fixed points of a recurrent neural network dynamics with binary neurons. We assume that the neural activation threshold is adjusted dynamically so that the population activity level maintains the same sparsity as the memories (see *Methods*, in subsection II H this assumption is modified). Synaptic dynamics are governed by three processes: Deterministic exponential synaptic decay [8, 11] (first term in eq.(1)), Hebbian learning [39] of new memories (second term in eq.(1)), and Hebbian consolidation of old memories following their reactivation (third term in eq.(1)),

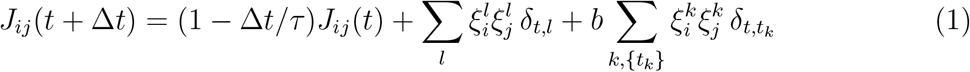

Here *J*_*ij*_(*t*) is the strength of the synapse between neurons *i* and *j* at time *t* (symmetric in *i* and *j*), 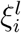 is the i-th element of the memory introduced first at time *t* = *l* and it is given by:

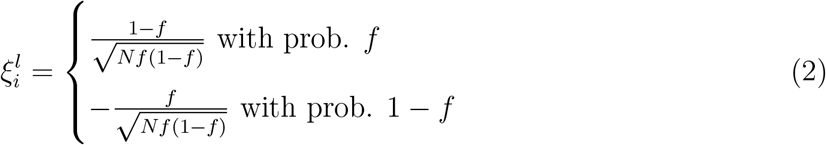

Here *f* is the fraction of neurons active in a memory state (the sparseness level). According to the above equation, new memories enter in each time interval Δ*t* and synapses decay at a rate 1*/τ*, representing the finite lifetime of synapses [40]. The last term represents a Hebbian strengthening of old memories following a sequence of reactivation events that occur for memory *k* at times denoted by *t*_*k*_ (which will be specified below). The factor *b* denotes the size of synaptic modification due to a single consolidation event of an old memory, assumed to be smaller than the Hebbian amplitude of learning a new memory (i.e. *b* < 1). The resulting connectivity matrix can be written as:

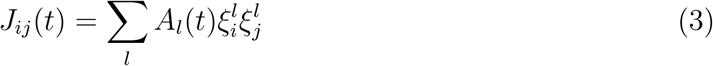

*A*_*l*_(*t*) is the *efficacy* of memory *l* at time *t*. The The ability to recall a memory depends on the level of noise, which originates from random interference with other memories. Its variance is proportional to the sum of the squares of all efficacies (see *Methods*):

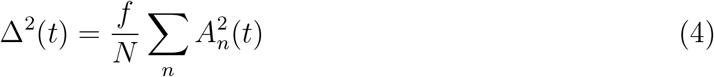

We define the critical efficacy *A*_*c*_, as the efficacy for which a memory pattern loses its stability. *A*_*c*_ is proportional to the interference noise Δ, with the proportionality constant depending only on the sparseness:

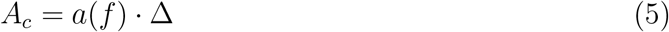

Where the factor *a*(*f*) can be approximated by (see Supplementary Information (SI) section 1):

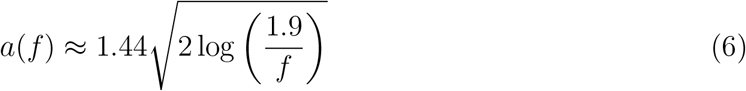

For *f* = 0.01 (which we will use throughout the paper), *a* ≈ 4.7.

### B. Pure forgetting

Without rehearsals, our model is similar to previous models of associative memory with forgetting [8, 10, 11], in which memory efficacies decay exponentially with age, *A*_*l*_(*t*) = exp(−(*t* − *l*)*/τ*) (Fig. 1a). Using eq.(4), the interference noise equals Δ^2^ ≈ *fτ/*(2*N*). For *τ* > *τ*_0_,where *τ*_0_ = 2*N/*(*fa*^2^(*f*)) (see eq.(5)), *A*_*c*_ increases above unity (the initial efficacy) and no memory will be retrievable. This *global catastrophic forgetting* is similar to the behavior of the Hopfield model after reaching memory capacity, where the interference effect is too strong and all memory states lose their stability [5, 6]. If *τ < τ*_0_, recent memories are retrievable, while memories older than a critical age 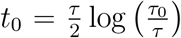 are forgotten (Fig. 1b). Thus, for short decay times, this model allows for continual learning of recent memories without global catastrophic forgetting. However, it predicts an unrealistic *age-dependent catastrophic forgetting*, where all memories up to a critical age are almost perfectly retrievable, and all older memories are completely forgotten. This sharp transition happens despite the graceful exponential decay of efficacies with age, and results from the collective effects of memory stability in the network.

**FIG. 1.**
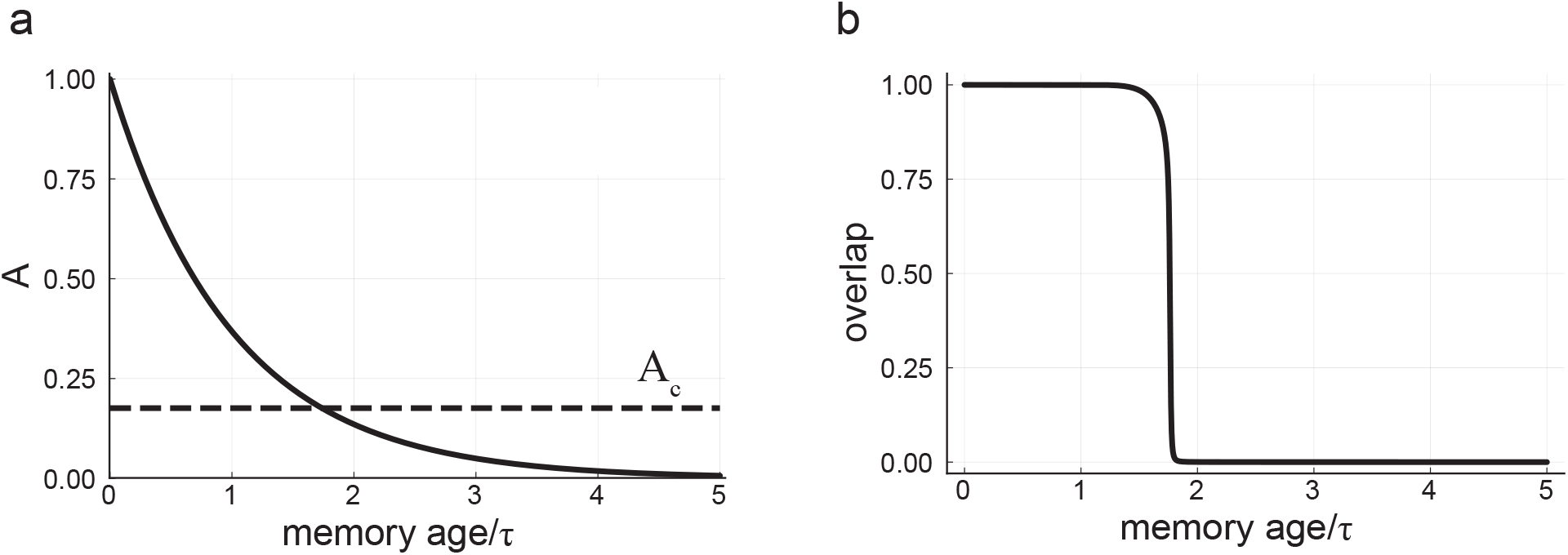
Pure forgetting. **a:** A memory efficacy trajectory as a function of time (solid line). The critical efficacy *A*_*c*_ is plotted as a dashed line. **b:** Overlap of the network state with a memory state as a function of the memory age. The overlap is a measure of memory retrievability - after initializing the network near a memory state, the overlap of the network activity with that memory after arriving to a steady state will be close to unity for retrievable memories and small compared to one for irretrievable memories. Here *N* = 8000, *f* = 0.01, *τ* = 2240. The catastrophic age here is ∼ 1.73*τ*, resulting in a capacity (number of retrievable memories) of 0.5*N*. Note the very large value of *τ* needed to support this capacity — this will be addressed in later sections.

In what follows we will show that when stochastic rehearsals are taken into account, the behavior changes dramatically, generating more realistic memory forgetting trajectories and allowing for lifelong memories.

### C. Nonlinear stochastic reactivation

To specify the statistics of reactivations, we revert to the continuous time version of Eq. (1) which yields the efficacies dynamics:

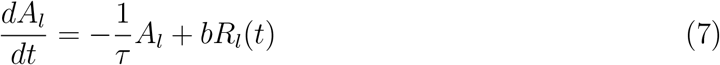

With *A*_*l*_(*t*) = 0 for *t < l* and *A*_*l*_(*l*) = 1. The reactivations are modeled as a point process

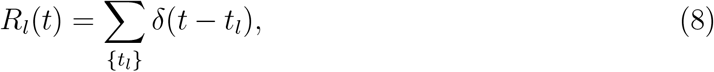

where *t*_*l*_ are the times at which memory *l* was rehearsed. To specify the rate of the reactivation process, we hypothesize that this process is more likely to yield a Hebbian strengthening of memories with large basin of attraction. The rationale is that during reactivation periods, the system is more likely to visit memories with large basins of attraction, stay there for a significant period of time triggering their Hebbian strengthening. In particular, memories that at some point in time lost their stability and are not attractors of the dynamics (i.e., have vanishing basin of attraction) will not be reactivated, will experience fast pure decay, and will be forgotten. Hence we model reactivation events as inhomogeneous Poisson processes, with mean rate *r*_*l*_(*t*) ≡ ⟨*R*_*l*_(*t*) ⟩

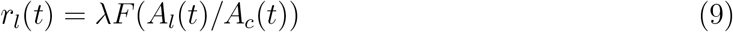

where *λ* denotes the maximal reactivation rate. As in eq.(4) and eq.(5), at all times *A*_*c*_(*t*) = *a*(*f*) · Δ(*t*). The nonlinear function *F* denotes the size of the basin of a memory and depends on the ratio of the memory efficacy over the critical capacity *A*_*c*_ (Fig. 2a). At any given time, only memories with non-zero basin size (i.e., *A*_*l*_(*t*) > *A*_*c*_ → *F* > 0) are retrievable and might be reactivated. Note that since the interference Δ(*t*) depends on the efficacies of all memories (eq.(4)), the reactivation rates of all memories are coupled in eq. (9) via *A*_*c*_.

**FIG. 2.**
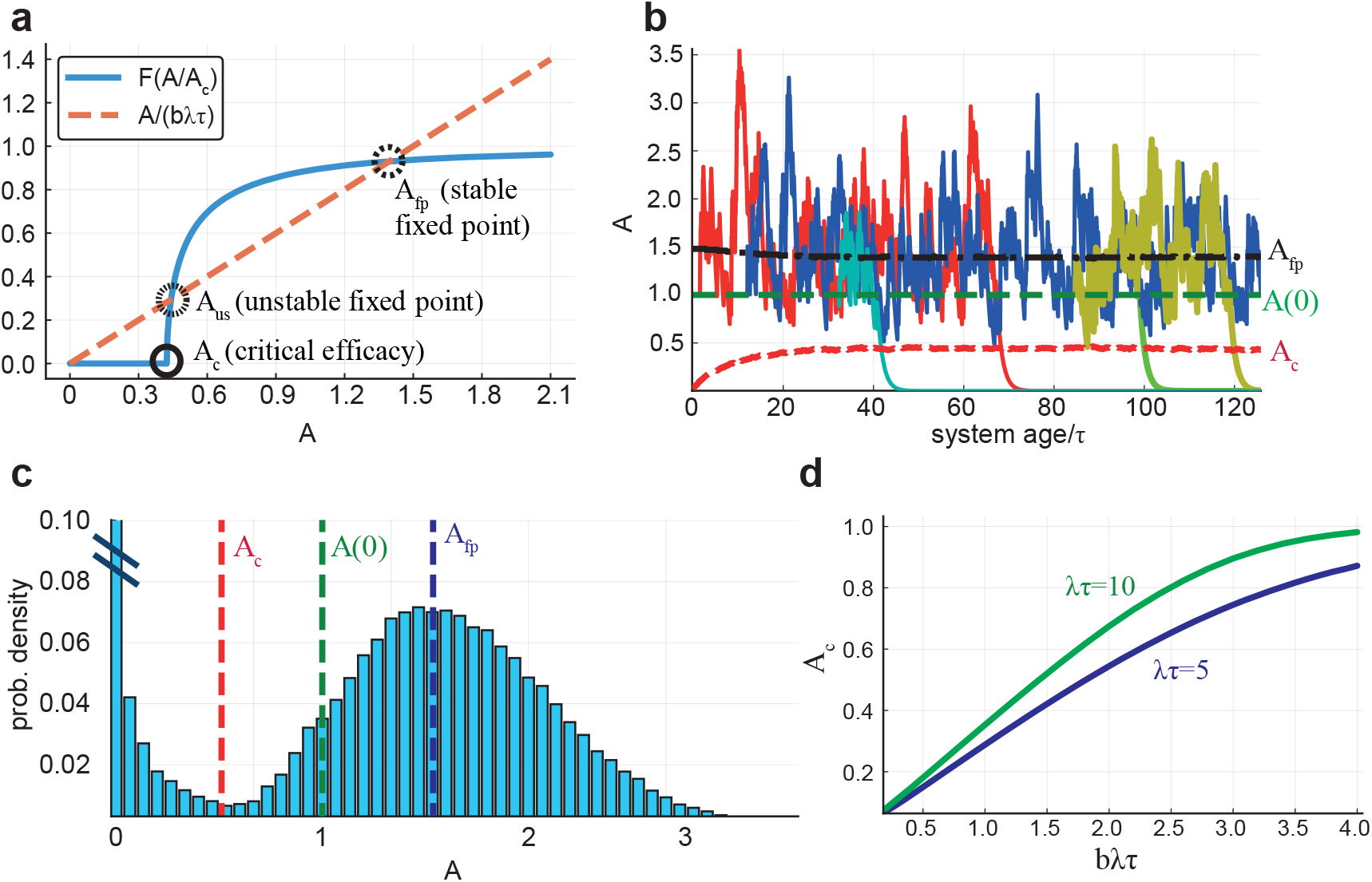
Stochastic memory dynamics.**a:** Blue: basin of attraction size *F* as a function of memory efficacy *A*. Orange dashed: the negative of the deterministic decay term in eq. (7). Importantly, *F* is zero for *A < A*_*c*_. Here *λτ* = 5, *A*_*c*_ = 0.4. **b:** Memory efficacies vs. age of the system. Memories enter with efficacy *A*(0) = 1, rehearsal efficacy *b* = 0.3. Most of them increase towards *A*_*fp*_ ≈ *bλτ* ≈ 1.5, and fluctuate around it. Large enough fluctuations can take efficacies below *A*_*c*_ (e.g, cyan curve at *age/τ* ≈ 40, yellow curve at *age/τ* ≈ 120). Some memories are alive for a very short time (e.g., green curve) and some for very long (e.g., red, blue curves). **c:** Distribution of memory efficacies after saturation of *A*_*c*_. **d:** Equilibrium values of *A*_*c*_ as a function of *bλτ* for different *λτ* values. Here *τ* = 160, *N* = 8000.

### D. The approach to steady state of memory consolidation

It is useful to first consider the average dynamics, replacing the reactivation point process by its mean rate, eq. (9). For a given *A*_*c*_, the resulting self-consistent equation for the steady state efficacies,

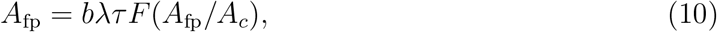

possesses two stable fixed points: one at zero and another one when the two competing processes, decay and reactivation, balance each other (*A*_fp_, Fig 2a). Due to the rapid saturation of the function *F*, for most of the parameter regime *A*_fp_ ∼ *bλτ*.

To fully understand the system’s behavior we need to consider the dynamics of *A*_*c*_ itself as well as the stochastic nature of the process. Initially, when the first memories enter the system, *A*_*c*_ ∝ Δ is very small and the memory efficacies consolidate around the value *A*_fp_ ∼ *bλτ*. As more memories are encoded, the interference grows and so does the critical efficacy (red line in Fig. 2b). When the critical efficacy is large enough, fluctuations in reactivation times lead some memory efficacies to drop below *A*_*c*_, making these memories irretrievable. A steady state is achieved when the flux of memories arriving at the system and consolidated is balanced by the rate of memories forgetting due to the drop of their efficacy below *A*_*c*_. At this stage, *A*_*c*_ reaches a fixed equilibrium value and so does the mean number of retrievable memories. The specific identity of the retrievable memories varies with time - some are forgotten while new ones are being consolidated. The distribution of efficacies at equilibrium (Fig. 2c) consists of two modes: The first is the contribution of the forgotten memories, below *A*_*c*_, which diverges at small *A* as *p*(*A*) = *τ/A*. The second, above *A*_*c*_, is a mode around *A*_fp_ representing the retrievable memories.

*A*_*c*_ increases as the amplitude *b* and number of reactivations per decay timescale *λτ* increase, due to increased interference (Fig. 2d). For moderate reactivation strength, *A*_*c*_ is well below both the encoding strength *A*(0) = 1 and the consolidation fixed point as seen in the examples in Figs. 2b,c. As reactivation strength grows, *A*_*c*_ increases and approaches 1, affecting adversely the consolidation process, as will be seen in the next section.

### E. The forgetting curve

Importantly, in our model, the time of forgetting of memories at a given age is highly variable, ranging from a fraction of the decay time *τ* (for unfortunate memories that weren’t rehearsed fast enough after learning), and up to hundreds of *τ* for well-rehearsed memories (Fig. 2b). Nevertheless, on average, memory retrievability decreases with memory age, and this is captured by the forgetting curve - the probability of retrieving a memory as a function of its age, after a steady state has been reached (Fig. 3a and Fig. 3b). This curve exhibits an exponential tail with a long time constant, denoted as the consolidation time *τ*_*c*_ - a direct result of the long time required for a large fluctuation in reactivation rates to form such that consolidated efficacies decrease from around *A*_fp_ to *A*_*c*_ (SI). The consolidation time enhancement factor *τ*_*c*_*/τ* can reach several orders of magnitude, allowing memories to survive for very long times compared to the intrinsic timescales of the system (as shown in Figs. 3c, d).

**FIG. 3.**
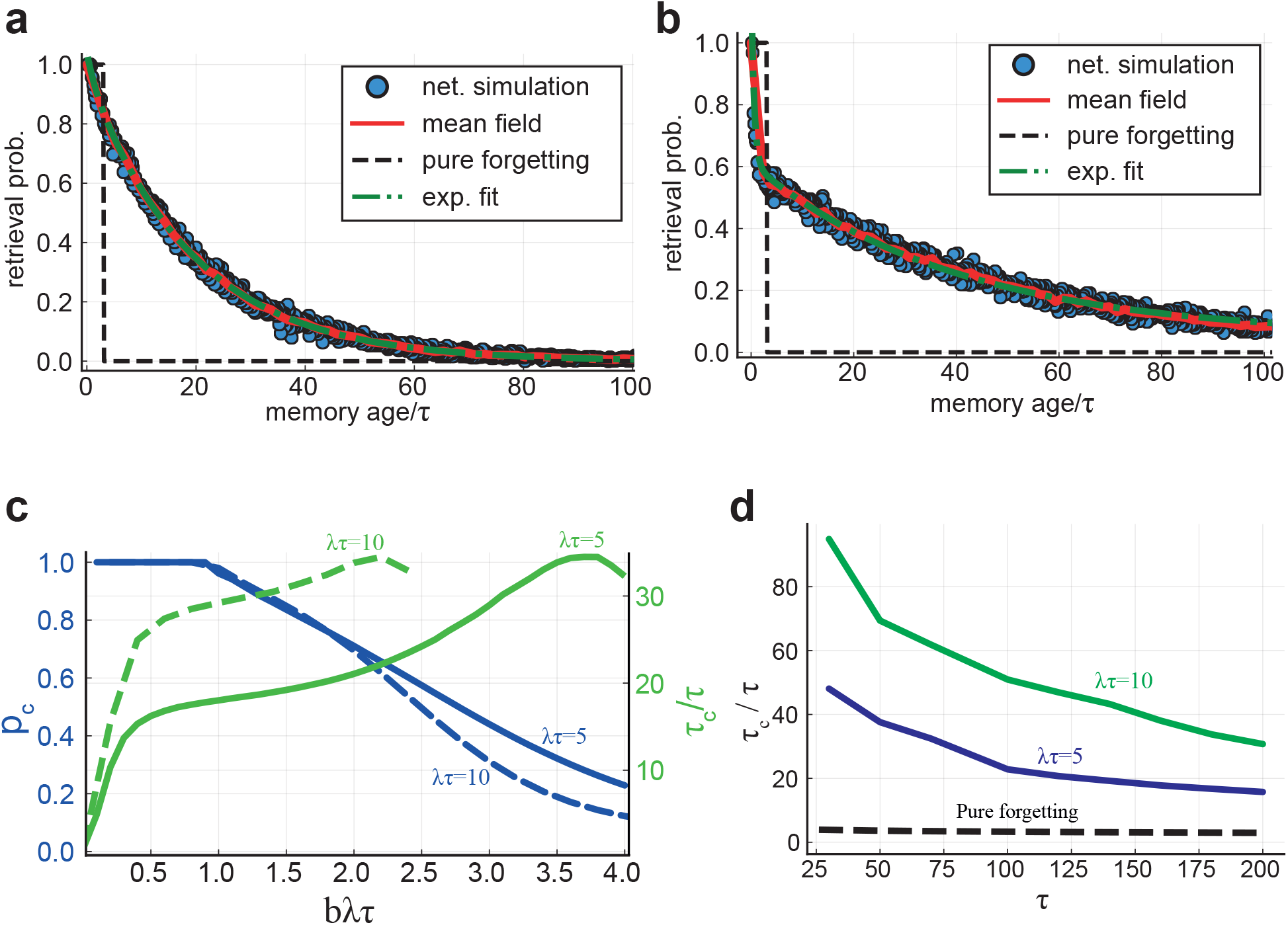
**a**,**b** *The forgetting curve*. The probability of retrieval as a function of memory age. Blue dots: full network simulation results (see Methods). Red solid lines: results of a mean field approximation (Methods). An exponential fit with characteristic decay time ≈ 18*τ* is shown in green (dash-dot line) in **a**, and a double exponential fit with characteristic decay times ≈ *τ* and ≈ 38*τ* in **b**. The retrieval probability for pure forgetting is shown in black (dashed line). In **a** *N* = 8000, *τ* = 160, *λτ* = 5, *b* = 0.3. In **b** same parameters as **a** except *b* = 0.25, *λτ* = 10. **c:** Blue (left y axis): Consolidation probability vs. *bλτ* for different *λτ* values. Green (right y axis): Consolidation time *τ*_*c*_ normalized by synaptic decay time *τ* vs. *bλτ* for different *λτ* values. Here *τ* = 160. **d:** Consolidation time *τ*_*c*_ normalized by synaptic decay time *τ* vs. *τ* for different *λτ* values. Blue curve: *λτ* = 5, *b* = 0.3. Green curve: *λτ* = 10, *b* = 0.25.

For fixed consolidation parameters *λτ* and *b*, consolidation time normalized by *τ* decreases with *τ* due to increased interference (Fig. 3d). For fixed *τ*, consolidation time increases sharply with *λτ* and *b* (Fig. 3c).

However, increasing these parameters may adversely affect the consolidation process. When *bλτ* is of order 1, most of the memories experience consolidation, as is the case in Fig. 3a. However when *bλτ* ≫ 1, *A*_*c*_ is close to the encoding efficacy (Fig. 2d). This causes a significant number of memories not to get consolidated. Therefore, the forgetting curve exhibits an initial fast decay with a characteristic time *τ*, in addition to the slow decay time *τ*_*c*_ (Fig. 3b). To quantify this effect, we measure the *consolidation probability* of memories, *p*_*c*_, defined as the chance of a memory efficacy to reach *A*_*fp*_, and therefore become part of long-lived memories. *p*_*c*_ decreases as a function of the reactivation strength from *p*_*c*_ = 1, for *bλτ <* 1, to zero for strong reactivation (Fig. 3d). The consolidation probability *p*_*c*_ together with *τ*_*c*_ are the key consolidation parameters.

### F. Capacity increases as power law with network size

We define the network’s memory capacity as the number of memories retrievable (memories with *A* > *A*_*c*_) in the equilibrium phase after long encoding time (Fig. 4). The capacity can be evaluated as the area under the forgetting curve. Hence, it can be approximated as (1 − *p*_*c*_)*τ* + *p*_*c*_*τ*_*c*_, where *p*_*c*_*τ*_*c*_ is the contribution from consolidated memories and fraction of memories that are consolidated and the first term is the contribution from unconsolidated memories. As seen previously, *τ*_*c*_ increases with *bλτ*, while *p*_*c*_ decreases with *bλτ* - less memories are consolidated, but the consolidated ones live longer. Maximal capacity is achieved when *bλτ* ≈ *A*(0) = 1, which is the maximal value that allows for 100% of the memories to get consolidated.

**FIG. 4.**
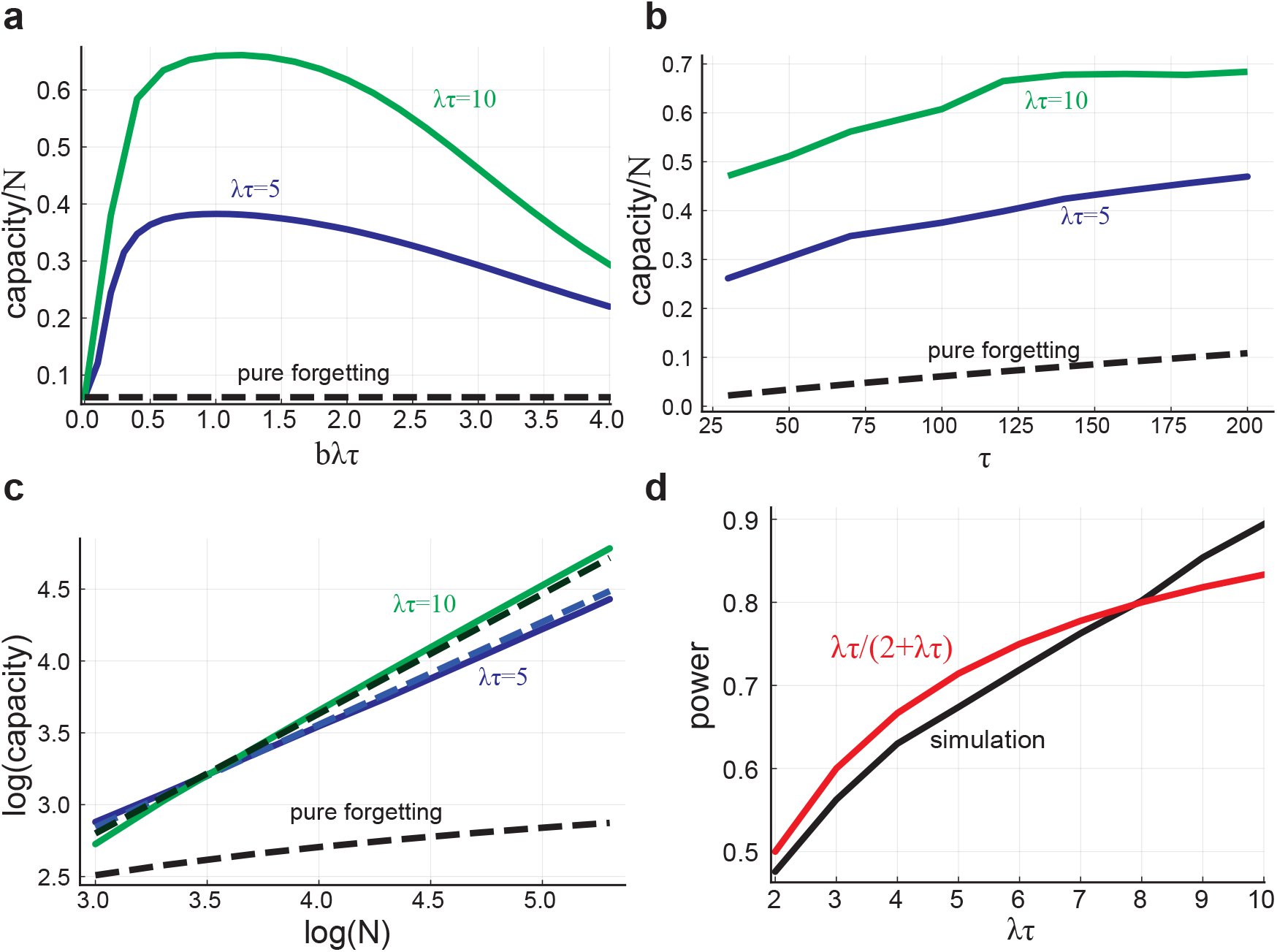
Memory capacity. **a:** The number of retrievable memories divided by *N* as a function of *bλτ* for different average number of rehearsals per characteristic decay time (*λτ*) values. The dashed line shows the capacity of the pure forgetting model. Here *N* = 8000, *τ* = 160. **b:** The number of retrievable memories divided by *N* as a function of *τ* for different *λτ* values. **c:** Capacity vs. N (logarithmic, base 10), solid lines show simulation results, dashed lines are analytical approximation. Here *τ* = 160, *b* = 0.3. **d:** The power of N vs. *λτ* (black), and the analytical approximation (red). Here *τ* = 160, *b* = 0.3.

To assess the efficiency of information storage in the network it is important to evaluate the dependence of the capacity on the network size, *N*. In previous ‘pure forgetting’ models [8, 10] the synaptic decay time was assumed to scale linearly with *N*, resulting in memory capacity *t*_0_ which is proportional to *N*. The same holds for our model. However, this scaling results in extremely large, biologically implausible, synaptic decay times for large networks. Here we assume that *τ* is a property of individual synapses and is independent of network size. Under this condition, the capacity in the pure forgetting model increases only logarithmically with *N*, Fig. 4c.

Interestingly, we find that in our model, the capacity scales as a power law of the number of neurons, with a power that approaches unity for large *λτ* values (Fig. 4c, d). To approximate the power analytically (for the parameter range where *p*_*c*_ ≈ 1), we first approximate *A*_*c*_, assuming that the main contribution to the interference noise comes from consolidated, retrievable memories (SI, sec.3):

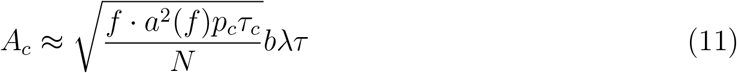

Now, under the assumption that *p*_*c*_ ≈ 1 and that the mean rehearsal rate is *λτ*, we get (SI):

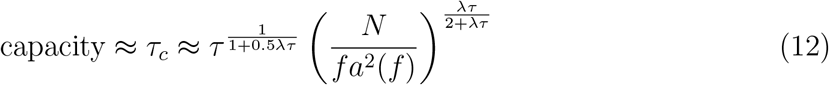

Figures 4c and d show the approximation gives a reasonable fit to the dependence of the capacity on *N*. Note the significant increase in capacity compared to the pure forgetting model.

### G. Inhomogeneity in initial memory encoding

So far we have assumed that all memories are encoded initially by Hebbian plasticity with the same amplitude *A*(0) = 1 (eq. 1). In reality, memories might differ in their encoding strength, for instance, due to factors such as attention, or emotional context. Thus, it is important to explore the effect of a distribution of initial encoding strengths. As long as most of the initial efficacies are in the neighborhood of *bλτ*, the global memory properties such as *A*_*c*_, forgetting curve, and capacity are not affected drastically. However, individual memories with initial efficacy below *A*_*c*_ are forgotten, while memories with *A*(0) larger than *bλτ* have slightly enhanced consolidation properties, as is confirmed in Fig. 5a for an exponential distribution of *A*(0) with mean 1.

**FIG. 5.**
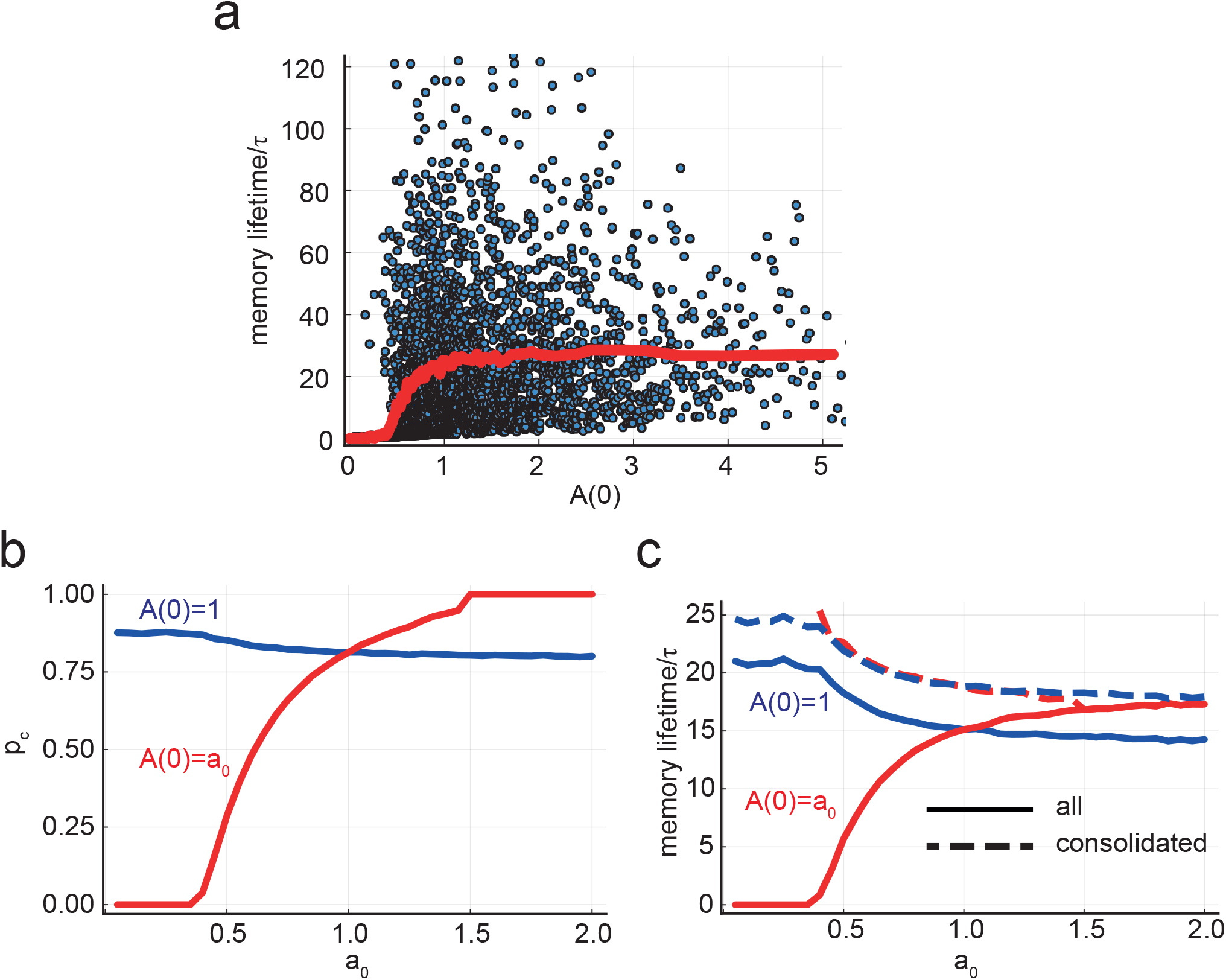
*Initial condition distribution*. **a:** Here *A*(0) for each memory is drawn from an exponential distribution with unity mean. Blue points are values for single memories, and the red line shows the mean. Note that the spread in lifetimes at each encoding strength is a result of the stochastic rehearsal process, which yields an exponential distribution of lifetimes and is present also in the uniform *A*(0) case. **b:** Consolidation probability vs. *a*_0_, which is the initial efficacy of half of the memories (the others have *A*(0) = 1). Values for memories introduced with *A*(0) = 1 are shown in blue, and for memories introduced with *A*(0) = *a*_0_ are shown in red. **c:** Memory mean lifetime (time from insertion to forgetting) as a function of *a*_0_. Same scenario and coloring as in **a**. Dashed lines are averaged lifetimes of *consolidated* memories only. Parameters: *N* = 8000, *f* = 0.01, *λτ* = 5, *b* = 0.3

To better elucidate the effect of inhomogeneity in *A*(0), we consider in Figs. 5b and c the case of a Bernoulli distribution, *A*(0) = *{*1, *a*_0_*}* with equal probability. For small *a*_0_ compared to *bλτ* = 1.5, the consolidation probability for memories with *A*(0) = *a*_0_ decreases drastically and vanishes for *a*_0_ below *A*_*c*_ ≈ 0.39 (Fig. 5b). When *a*_0_ increases above 1, consolidation probability of these memories increases until it reaches 1 for *a*_0_ ≈ *bλτ*. On the other hand, memories with *A*(0) = 1 are only moderately affected by changing *a*_0_. The mean lifetime of memories with *A*(0) = *a*_0_ *<* 1 drops considerably (Figure 5c and a). This is, however, due to averaging the lifetime of all memories including those that did not consolidate. Importantly, in our model, memories with small *a*_0_ that did reach the neighborhood of the fixed point have the same long lifetime as other consolidated memories, independent of the original encoding strengths as shown by the dashed lines in Fig. 5c.

### H. The effects of structural perturbations on memory function

In this section we analyze the effect of damage to the circuit on memory storage and retrieval. In previous sections, we assumed for simplicity that the neuronal firing threshold is automatically adjusted to guarantee a fixed mean activation level, *f* (see Methods). Here we assume that the firing threshold is fixed since we anticipate that part of the effect of perturbation is the disruption of the level of activity. Importantly, in the case of constant threshold, the dependence of *A*_*c*_ on Δ is not linear. It has a non-zero value for small Δ reflecting the requirement for the encoding efficacy to be large enough for neurons to cross the threshold. Above some critical Δ*, A*_*c*_ rises abruptly, causing all memories to lose stability, due to over-activation of the network when the noise level is high (Fig. S4). At equilibrium Δ is below but close to the critical value (for the presented parameter range). In this scenario, the properties of the unperturbed system are similar to those of the fixed activity scenario, with a memory capacity that depends on the threshold value. For the presented results we used the threshold value 0.36 which maximizes capacity in unperturbed conditions (See SI).

#### 1. Noisy synaptic dynamics

We first consider perturbations of the synaptic learning and consolidation processes by adding white noise *χ* to the synaptic dynamics, for all *t* ≥ *t*_onset_,with a diffusion coefficient *D*,

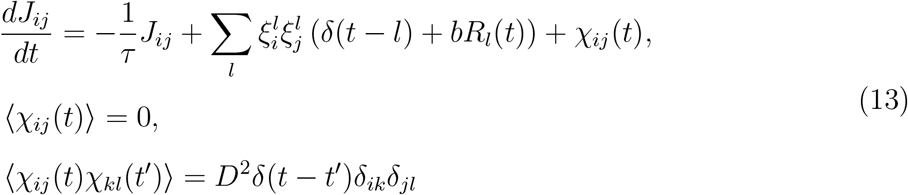

The effect of this noise is approximately an additive contribution to the total variance of local fields

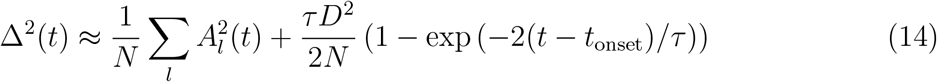

After the noise onset, Δ increases rapidly above its critical value, causing a sharp increase in *A*_*c*_, and the blocking of rehearsals for all memories. This in turn causes a rapid decrease in magnitude of stored memory efficacies, leading to a decrease in Δ below the critical value and a decrease in *A*_*c*_ to a value which is between the value before the onset and the value just after the noise onset. This new equilibrium value of *A*_*c*_, with reduced capacity, occurs over ∼ *τ* (Fig. 6a). Although overall reduction in capacity may be mild for moderate *D* values, there is a large reduction in the retrieval probability of memories that were encoded around the perturbation onset time, due to the sharp transient increase in *A*_*c*_. In contrast, memories that have already been consolidated suffer only a mild reduction in survival probability (relative to unperturbed memories of the same age). Likewise, newly entered memories have s high probability of consolidation, since they experience the equilibrium value of *A*_*c*_ and their retrieval probability is similar to the unperturbed case (Fig. 6b).

**FIG. 6.**
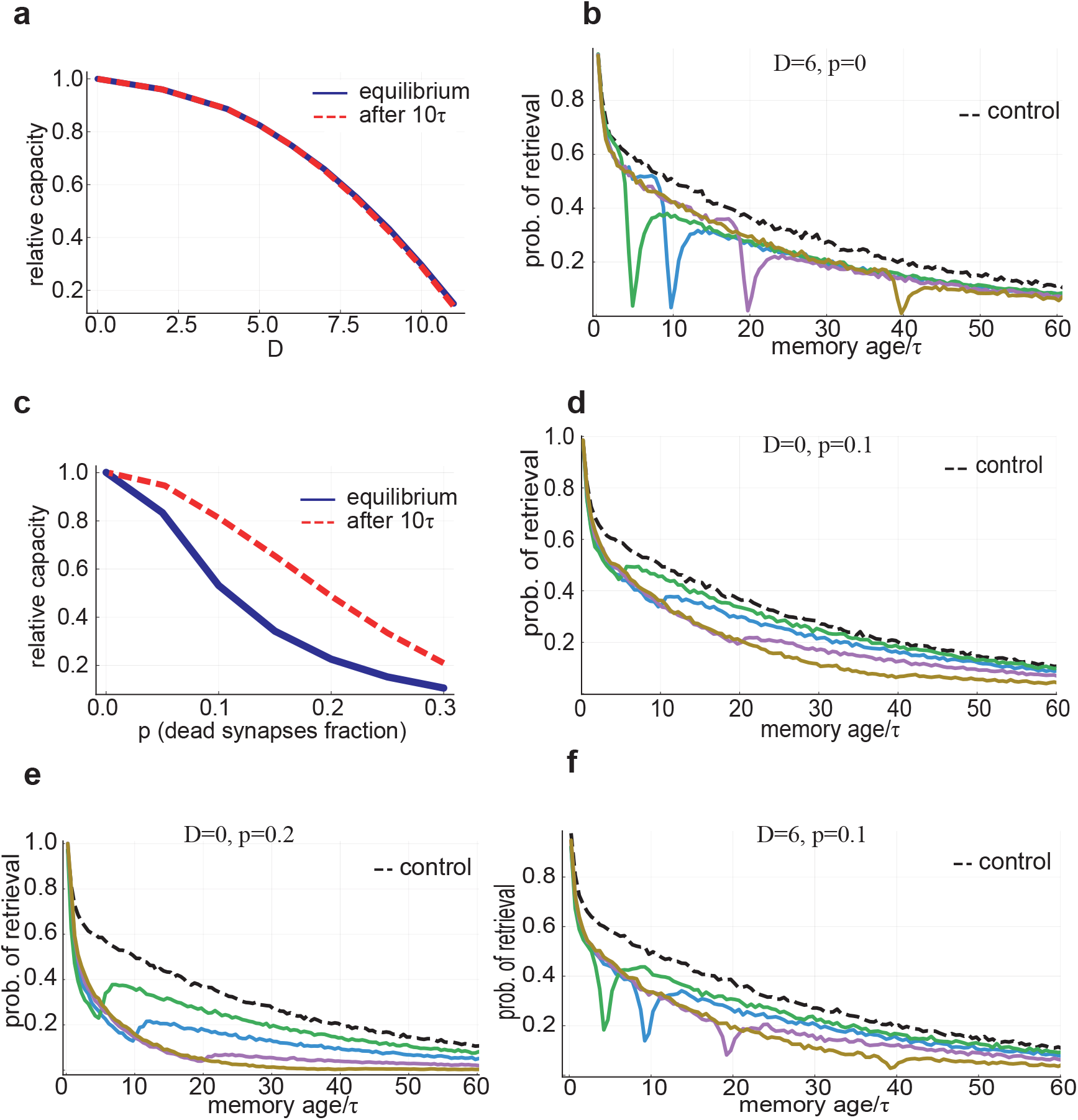
Perturbations and memory deficits. **a:** The ratio between the capacity with and without injected noise vs. the diffusion coefficient *D*. **b:** Retrieval probability vs. memory age with noisy synaptic dynamics (*D* = 6). Noise onset was before: 5*τ* (green), 10*τ* (blue), 20*τ* (purple), 40*τ* (brown). The control (black) is simulated with noiseless dynamics. **c:** The ratio between the capacity with and without synaptic dilution vs. the silenced synapses fraction *p*. **d:** Retrieval probability vs. memory age for random synaptic dilution (*p* = 0.1). Coloring as in **b. e:** Same as **c**, but with *p* = 0.2. Memories of all ages are affected, with some non-monotonicity caused by the small efficacies of newly learned memories, dropping more easily below *A*_*c*_. **f:** Combination of synaptic dilution and noisy synaptic dynamics, *D* = 6 and *p* = 0.1. Coloring as in **b**. Parameters: *N* = 8000, *τ* = 160, *λτ* = 10, *b* = 0.25

#### 2. Random synaptic silencing

Another perturbation we consider is the death of a fraction of the synapses. We model the effect of the synaptic death by multiplying the connectivity matrix *J*_*ij*_ by a binary random dilution *{*0, 1*}* matrix:

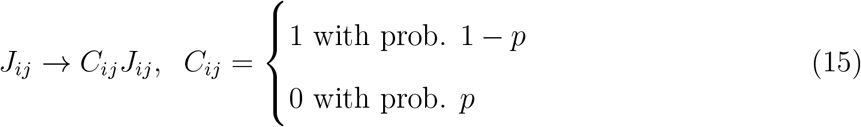

Unlike the additive noise considered above, synaptic dilution process is multiplicative, reducing both the effective efficacy of each memory (by a factor 1 − *p*), and the interference noise Δ (by factor 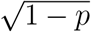), and in general reduces the signal to noise ratio (Methods).

After the dilution onset, *A*_*c*_ barely changes (due to the weak dependence of *A*_*c*_ on Δ in the constant threshold scenario), while all the efficacies are reduced, causing a reduction of retrievability that spreads over the entire age range (Fig. 6c and d) and a new equilibrium is achieved slowly.

Interestingly, neural adaptation (modeled here as a decrease in the neural activation threshold) can reduce the memory loss due to silencing (also reported in [41–43]), by reducing the minimum efficacy required for activation, i.e., *A*_*c*_, thereby recovering some of the gap between memory efficacies and *A*_*c*_ (Fig. 7).

**FIG. 7.**
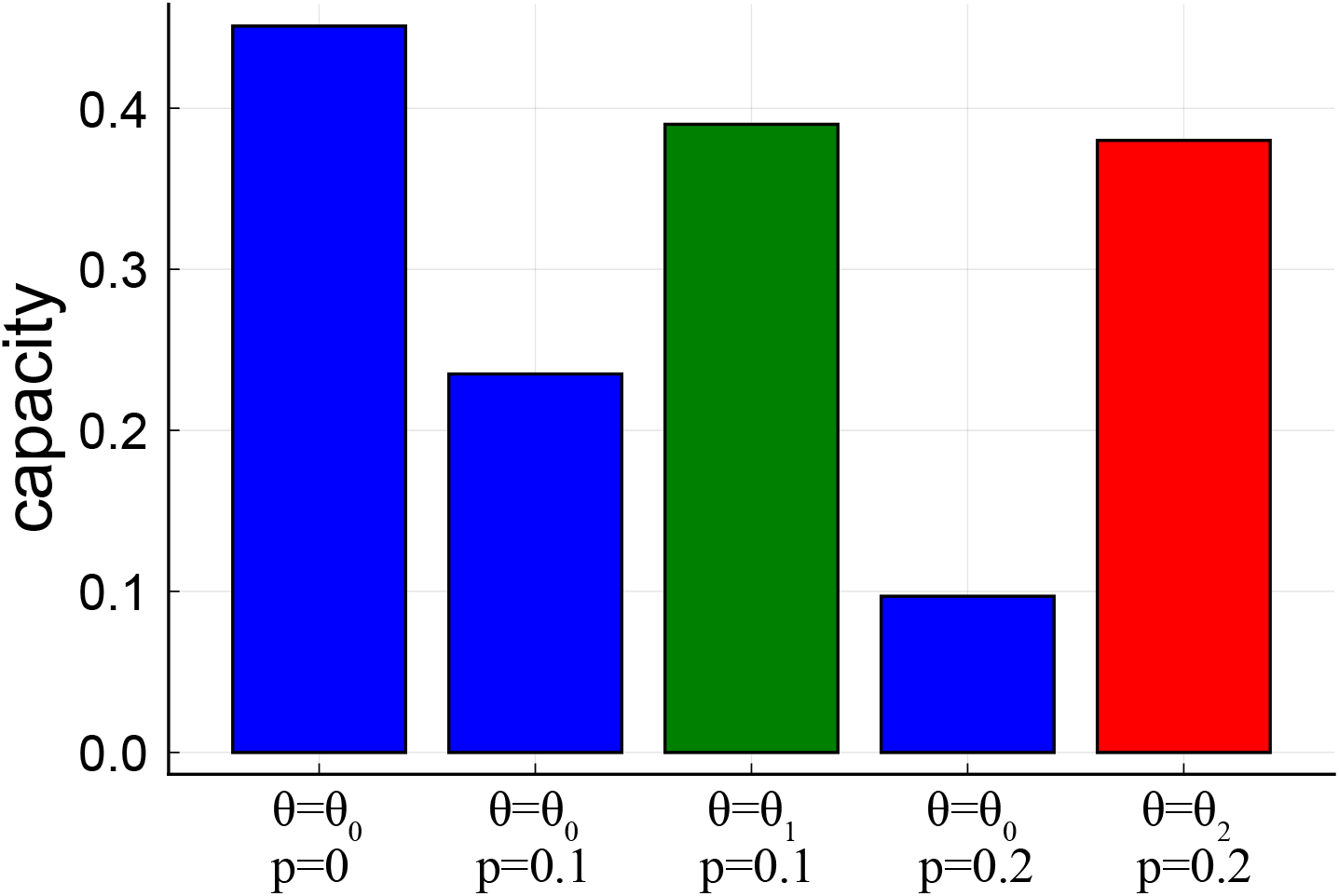
Effect of threshold adaptation. Blue bars show capacity with threshold optimized for the noiseless case (*θ*_0_ = 0.36). Green bar shows capacity with threshold optimized for low dilution (p=0.1, *θ*_1_ = 0.31). Red bar shows capacity for threshold optimized for high dilution (p=0.2, *θ*_2_ = 0.29). Parameters: *N* = 8000, *τ* = 160, *λτ* = 10, *b* = 0.25

Thus, our model predicts a qualitative difference in the effects of the two types of perturbations: synaptic dilution affects memories of all ages, causing a reduction in capacity that develops over a long time and can be partially compensated for by threshold adaptation, while additive synaptic noise results in a deficit largely confined to the time of the perturbation onset, and fast convergence to a new equilibrium.

### I. Distribution of synaptic decay times

Experiments showing that the time scale of synaptic and spine turnover is variable [40], and observations of power law memory retention curves in some memory studies [1–4, 44] encourage consideration of the properties of synaptic dynamics with heterogeneous decay time constants, yielding,

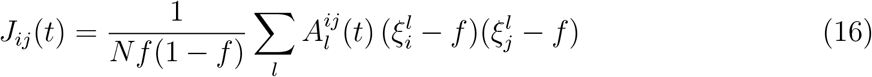

where the efficacies 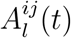 contributed by each synapse obey

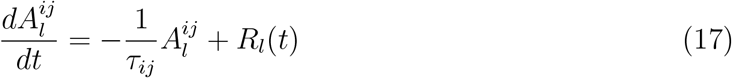

The mean efficacy of each memory is the average over these contributions,

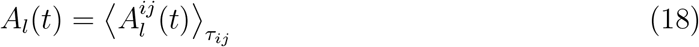

where ⟨..⟩_*τij*_ denotes the average over the distribution of synaptic time constants. Likewise, the noise term is proportional to the sum of second moments of the efficacies:

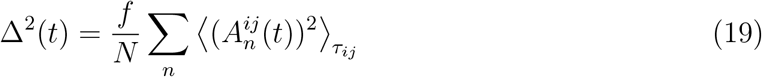

As an example, we show the case where the decay time constants are power law (Pareto) distributed, i.e., *P* (*τ*) ∝ *τ* ^−(*α*+1)^ ; *τ* ≥ *τ*_0_, *α* > 0 (Methods). In the absence of rehearsals (pure decay), there will be a global catastrophic forgetting for *α* ≤ 1, where the mean of the decay rate, and therefore the interference noise, diverges. For *α* > 1 there will be a catastrophic age dependent forgetting, as in the case with a uniform decay time scale (See [13] SI). With stochastic nonlinear rehearsals, for large *p* the forgetting curve is approximately exponential, similar to the single *τ* case (Fig. 8b). This is because the dominant contribution comes from the shortest time *τ*_0_. Interestingly, for intermediate values (1 *< α <* 1.8), the forgetting curves have an approximately power-law decay (Fig. 8a). In this regime, the retrieval probability is affected by contributions from a broad range of time constants: neither the minimal *τ* nor outliers with very large values are dominant.

**FIG. 8.**
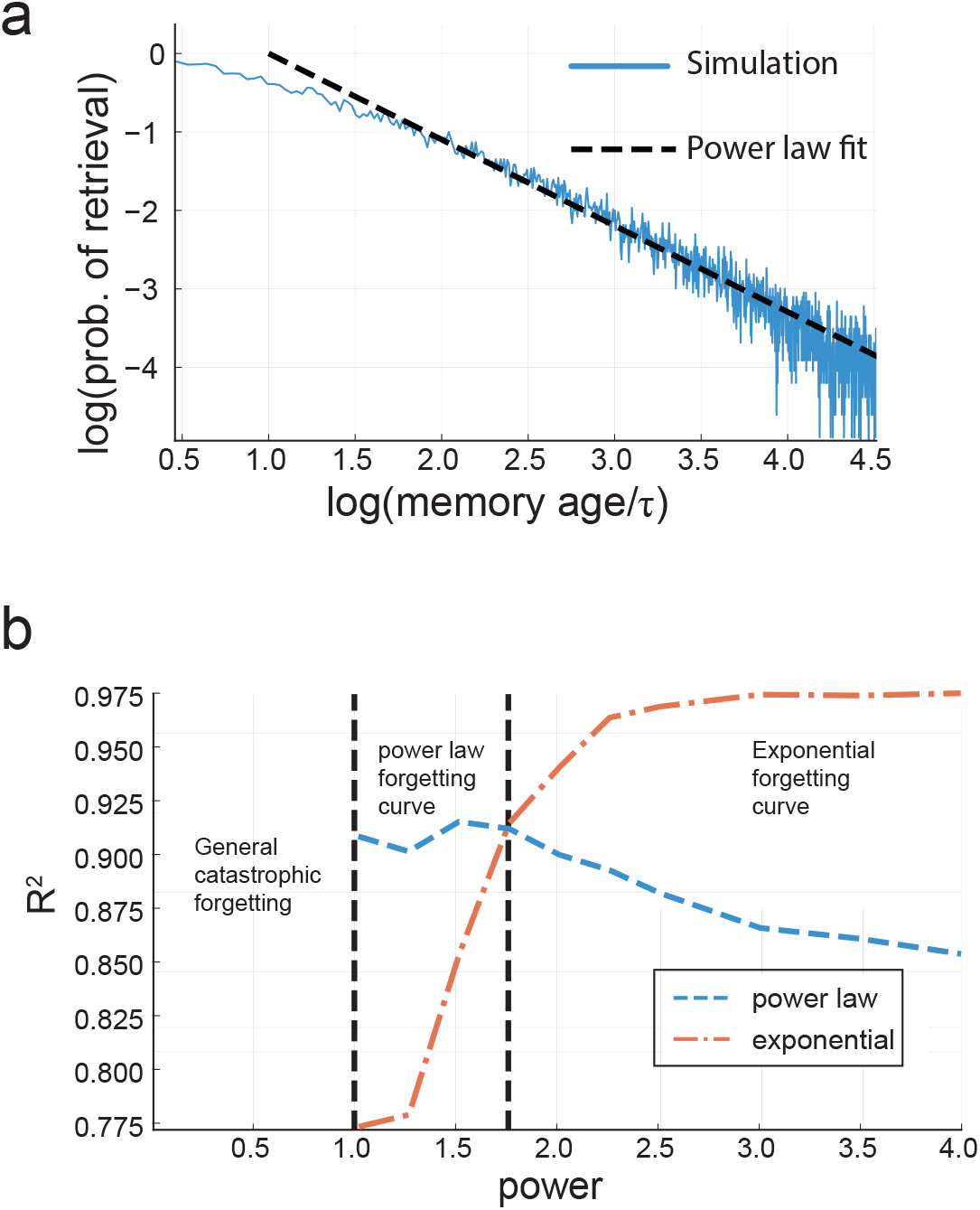
Power-law synaptic decay characteristic time distribution. **a**: Retrieval probability as a function of memory age on log-log scale (blue), with power-law fit in black (dashed line, slope≈ 1). Here *λτ* = 5 for the empirical mean *τ, b* = 0.25, the power *α* = 1.5. The minimal *τ* is 20. **b**: Goodness of fit (*R*^2^) for the forgetting curve using an exponential function (orange, dash-dot) and power-law function (blue), as a function of the power parameter of the characteristic time distribution.

## III. DISCUSSION

### A. Consolidation time scale

We have proposed a stochastic self-amplified memory consolidation mechanism and showed that it leads to smooth forgetting curves that extend much longer than the synaptic decay time. Our model provides estimates for the global long-term memory properties such as the capacity of the network, the shape of the forgetting curve and the average lifetime of memories. Translating synaptic decay time to realistic times is hard. In rodents, spine turnover time is estimated to be of the order of several weeks in the hippocampus and up to a year in the cortex [40, 45–48]. In humans, these times may be longer given the lower metabolic rate [49, 50]; however, there are no direct experimental data. In addition, the model synaptic decay time *τ* is in units of the mean inverse rate of encoding of episodic memory, which is hard to estimate, but it is likely to be of the order of weeks. Thus, assuming a human spine turnover time of the order of months yields *τ* of the order of tens of months, which could lead to mean memory lifetime of several years. At present, these estimates are speculative.

### B. Memory deficiencies

We have considered two types of perturbations to the memory circuit: synaptic death and increased synaptic noise. Both types of damage result in reduced retrievability of memories introduced prior to the damage onset, a phenomenon known in the literature as retrograde amnesia [14–17]. Due to the consolidation effect in our model, the amnesia is temporally graded: memories learned just before the noise onset are more severely affected than older ones, because they didn’t have enough time to consolidate, and were more fragile at the onset time. This effect is more prominent in the case of increased synaptic noise than synaptic dilution, due to the sharper drop in basins of attraction size after the noise addition. Perturbations cause a drop in retrieval performance of new memories entering after the perturbation onset, a manifestation of anterograde amnesia. This effect is temporally graded as well, being more severe for memories introduced just after the onset, and is especially prominent deficit in the additive noise case. Another interesting difference is the approach to a new equilibrium, which is fast in the case of additive noise but slow in the dilution. In addition, threshold adaptation can compensate part of the memory dysfunction caused by dilution, but not by additive noise.

Our model also allows for exploration of transient perturbations (SI), where the damage lasts for a finite time window [51]. In this case there is again a temporally-graded retrograde amnesia. Interestingly, new memories introduced after the end of the event not only regain retrievability, but can even improve their retrievability compared to control. This is due to the increased forgetting rate during the event, which results in lower interference noise and increased rehearsal rate after the event end.

The predictions of our model should be contrasted with the pure decay model where similar perturbations reduce the capacity (maximal age for retrievable memories), but don’t introduce any non-monotonicity in the forgetting curve, which is still a step function, but with a reduced width.

### C. Relation to previous models

In the classical theory of systems memory consolidation [14–16, 18], the interaction between the hippocampus (HC) and the cortex plays a central role, with HC storing memories for short period of time, and following rehearsals, memories are transmitted to the cortex for long-term storage. In the recent Multiple Trace Theory (MTT) [18, 27] autobiographic memories are stored for long term memory in both HC and cortex and consolidated through rehearsals that establish multiple memory traces in HC. This model shares some key elements with our theory, such as ongoing, life-long consolidation of memories and rehearsals which make memories more robust to perturbations. However, it is unclear how MMT can scale to large numbers of stored memories. In addition, in [27] the rehearsal statistics (new trace formation) don’t depend on the robustness of the memories, nor the model takes into account interference between memories.

A few studies used neural network models where rehearsals are modeled as random visits of learned memories [29, 31, 34], or implicit rehearsals (via memory traces embedded in the noise correlations [28]). Yet, none of these papers demonstrated consolidation in a large network with a number of consolidated memories scaling with network size. They also consider rehearsals of a finite batch of previously encoded memories rather than with life-long learning. Benna and Fusi [13] studied memory storage with complex synapses, where a consolidation process is implemented in the dynamics of synapses, with a cascade of synaptic characteristic times. They show that their mechanism gives rise to a power-law decay of the signal-to-noise ratio (SNR, equivalent to *A*_*l*_(*t*)*/*Δ(*t*) in our model) with age. However, this model still exhibits a deterministic catastrophic age-dependent forgetting, such that all memories older than a critical age are non-retrievable, whereas all newer memories are almost perfectly retrievable. A recent phenomenological model [44] derives a power-law form for memory retention curves with a power of 1 or smaller. A power close to 1 for intermediate ages is consistent with our result for a distribution of synaptic decay time. However, at present, it is unclear whether the experimental paradigms and time scales in which a power law is observed are relevant to life long episodic memory.

Fiebig and Lansner [33] proposed a three component model, each with different synaptic decay rate, which performs continual learning with self-generated rehearsals. Similar to our work, they study the effects of perturbations and show similarities to human data. However, this work does not provide an analysis of the model, and does not explore the dependence on the different parameters such as network size and synaptic decay time and rehearsal rates. Comparison with our results is hampered also by the fact that synaptic decay in their model is an active process, dependent on memory arrivals among other factors.

### D. Limitations and future work

In this paper we don’t explicitly model the rehearsals process - how the system moves between activation states and visits different attractors. Possible mechanisms could be destabilization of attractors by adaptation [34, 52] or transitions induced by random initialization processes [29]. These mechanisms will generate a rate of rehearsals per memory that depends on its basin of attraction size, as in our model, but whether the rate is simply proportional to the basin’s size as we assume is yet to be tested.

Our model can be extended in a variety of ways, including more biologically plausible neuronal and synaptic integration. Additionally, our framework allows for analyzing the effect of other types of perturbations, such as post-traumatic stress disorder amnesia [53, 54].

### E. Conclusions

The stochastic nonlinear rehearsal mechanism proposed in our work is, to the best of our knowledge, the first large-scale memory model that gives rise to realistic gracefully decaying forgetting probability curves, with exponential or power law tails depending on synaptic decay rate distribution. Our model’s capacity scales as a power law of the number of neurons, with a power that approaches unity for a large mean number of rehearsal events per synaptic decay time. Our model predicts that the onset of perturbation to the circuit, in the form of synaptic noise, leads to non-monotonic memory deficits affecting more strongly memories encoded around perturbation onset time, which have not yet a chance to consolidate. The richness of the model behavior in normal and diseased conditions provides a theoretical framework for predictions and testing against empirical data on human memory.

## IV. ACKNOWLEDGEMENTS

Research of NS was partially supported by the Swartz foundation and by the Center for Brains, Minds and Machines. HS is supported in part by the Gatsby Charitable Foundation. GK and HS are supported by NIH and NSF (the CBMM Center).

## V. METHODS

### A. Network model

As described in section II, memories are modeled as sparse, uncorrelated *N* dimensional activation patterns (N is the number of neurons), and the synaptic dynamics are governed by three processes: deterministic synaptic decay with rate 1*/τ*, Hebbian learning of new memories, and rehearsal of old memories, which is the central novelty of our model. In continuous time, eq. (1) becomes:

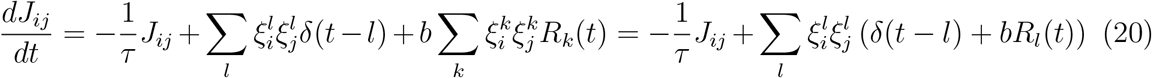

The rehearsals are modeled as a point process

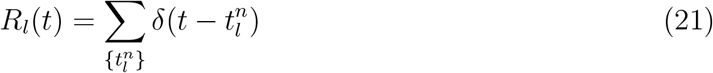

Inserting the ansatz (3), we get that *A*_*l*_ obeys the differential equation:

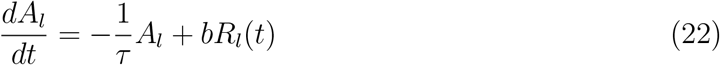

With *A*_*l*_(*t*) = 0 for *t < l* and *A*_*l*_(*l*) = 1.

The single neuron dynamics are binary, and given by:

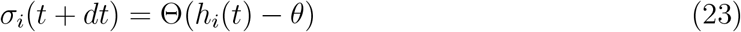

where *σ*_(_*t*) is the state of neuron *i* at time *t*, Θ(*x*) is the Heaviside step function, *h*_*i*_(*t*) is the local field (total input received by neuron *i* at time *t*):

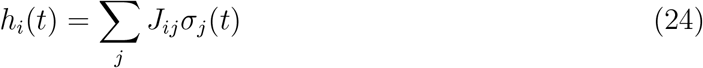

and *θ* is a threshold, set at every time step such that the total activation of the network is maintained and equal to *f N* (practically, in full simulations, at each time step we choose the *f N* neurons with the largest local fields and set their state to one, and all the others are set to zero).

#### 1. Mean field equations, basins of attraction

We would like to find the relation between memory stability, measured by the memory pattern’s basin of attraction size, and the efficacy of the memory and all other memories in the system. First, we define two useful quantities: 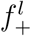 is the probability of a neuron to be active in the current state given that it is active in the memory pattern 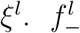 is the probability of a neuron to be active in the current state given that it is not active in the memory pattern *ξ*^*l*^. In other words, 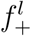 is the fraction out of the neurons active in memory state *l* that are active in the current state. 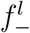 is the fraction out of the neurons not active in memory state *l* that are active in the current state. We will omit the *l* dependence of *f*_*±*_ from now on. In terms of these quantities,

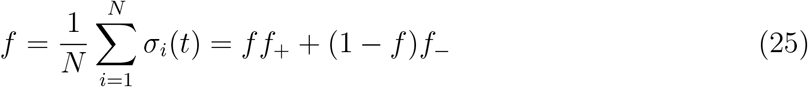

For clarity, in this section instead of the memory patterns definition we use above (eq. (2)) we define the patterns in an equivalent, more explicit way:

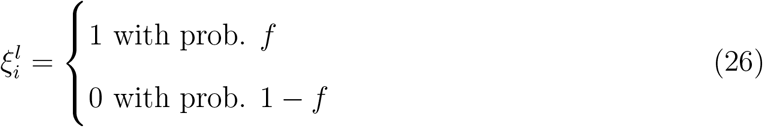

, and we normalize the connectivity matrix accordingly.

The overlap between memory pattern *l* and the system’s state *σ*

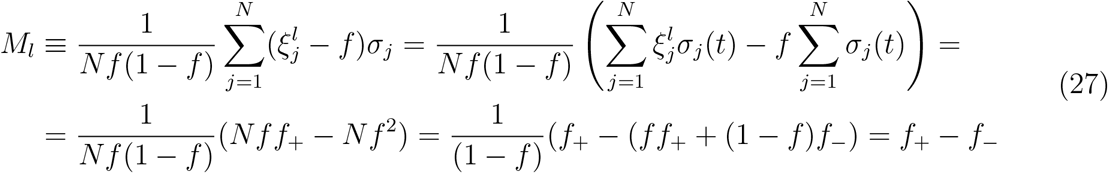

Now, assume that the current state is close to the memory state *ξ*^*l*^. The input to neuron *i* which is active in memory pattern 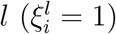:

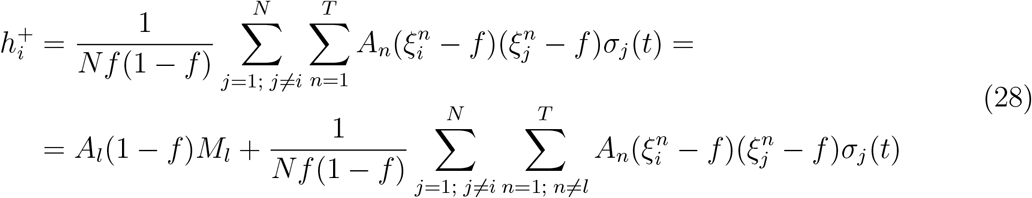

Averaging 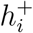 over memories realizations gives *A*_*l*_(1 − *f*)*M*_*l*_, and the variance:

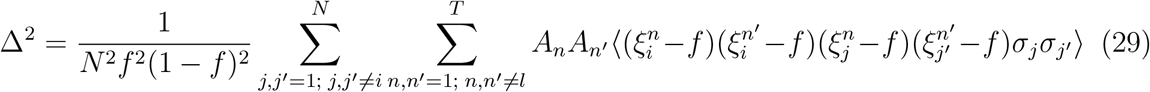

Now, because the system state is close to the memory state *ξ*^*l*^, we can assume it is uncorrelated with all other memory states. Hence, there are contributions only from terms with *j* = *j′, n* = *n′*:

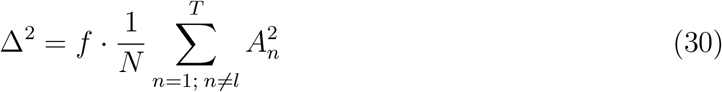

Applying the central limit theorem, we approximate 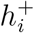 by a Gaussian variable with mean *A*_*l*_(1 − *f*)*M*_*l*_ and variance Δ^2^. Now, *f*_+_ is the probability for 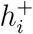 to be larger than *θ*, which is given by the complimentary error function, 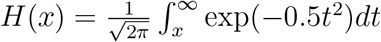:

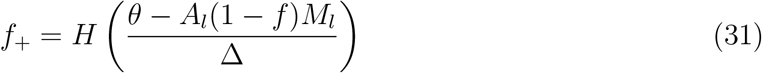

In a similar way (same noise term, mean equals −*A*_*l*_*M*_*l*_*f*) we find for *f*_−_ (for *f ≪* 1) :

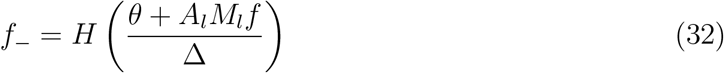

Note that we didn’t set the threshold *θ*, but instead demanded a constant population activation *f*. Equations (25), (27), (31) and (32) allow us to write an equation for the overlap dynamics:

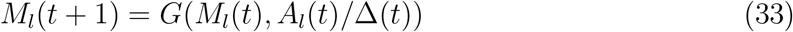

where

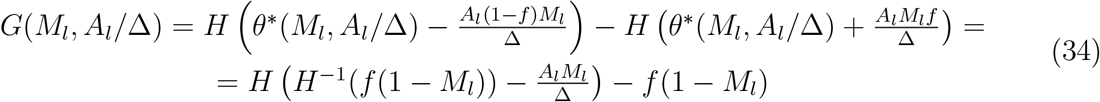

and

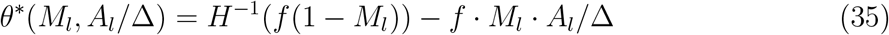

Therefore, the equation for the overlap fixed points is:

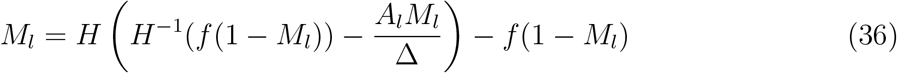

Now, we numerically find the fixed points for *M* at a given *A/*Δ by running the dynamics described by eq. (33). Typically (for large enough *A/*Δ), there will be a stable fixed point at *M* = 0, a stable fixed point 0 *< M*_*s*_ ≤ 1 and an unstable fixed point 0 *< M*_*us*_ *< M*_*s*_. We approximate the basin of attraction size as the distance *M*_*s*_ − *M*_*us*_. This way, we obtain the basin size as a function of *A/*Δ, *F* (*A/*Δ). We check the validity of our approximations by simulating a full neural network model and checking numerically the basin of attraction sizes, and find good agreement (SI), which is the basis for the good agreement in retention curves between the mean field simulations and the full network simulation (Fig. 3a,b). We define the critical efficacy *A*_*c*_ as the efficacy for which *M*_*s*_ = *M*_*us*_ (meaning, the non-zero overlap solution loses stability). This happens approximately when *M*_*s*_ = *M*_*us*_ ≈ 0.85. As one can see from equation (36), the fixed points depends only on the ratio *A/*Δ and on *f*, and therefore *A*_*c*_*/*Δ is only a function of *f*, and we can write *A*_*c*_ = *a*(*f*)Δ. For *f* = 0.01 (the typical value we use throughout the manuscript) we find numerically that *a*(*f*) ≈ 4.7. Analytical approximation for *a*(*f*) is given in the SI.

### B. Numerical simulations

In our simulations, we first numerically solve the coupled stochastic differential equations for the efficacies (Eq. (7)). Theoretically our model considers infinite number of memories. However, practically we solve the equations for a finite but large number of memory efficacies, typically 200*τ* − 1000*τ*, chosen such that *A*_*c*_ saturates to its steady state value. We measure time in units of the lag between the introduction of two consecutive memories (assumed constant). At every integration time step *dt* (small compared to all characteristic timescales of the system, typically *dt* = 0.05*/λ*), a rehearsal event might occur for each memory with *A*_*l*_ ≥ *A*_*c*_. We generate a uniform random number between 0 and 1 and compare it to *λ · F* (*A*_*l*_(*t*)*/*Δ(*t*) *· dt*. A rehearsal event of memory *l* will happen at time *t* if the uniform random number is smaller than *λ · F* (*A*_*l*_(*t*)*/*Δ(*t*) *· dt*. This approximates the statistics of a non-homogeneous Poisson process. By averaging over many such realizations (typically 500), we calculate the efficacy histogram, capacity (counting how many efficacies are above *A*_*c*_) and retrieval probabilities (by checking the probability for the efficacy of a memory introduced *l* time units into the past to be retrievable now). These calculations are referred to as “mean field simulations”, and they don’t include generation of random memories and building the connectivity matrix.

#### Full network simulation

When simulating the full network model, after generating the efficacies, we randomly generate memory patterns (binary vectors of dimension *N*) to be stored, and build the connectivity matrix according to eq. (3). Then, to measure retrievability, we initialize the network’s state at a memory pattern, and let the binary neurons dynamics run until they settle to a steady state. Then, we measure the overlap between the pattern and the steady-state activity. We say a memory is retrievable if the overlap is ≥ 0.85. For measuring the basin of attraction sizes of the memory patterns, we generate an initial state by randomly flipping the state of units in the memory pattern (conserving the total activation *f N*), and run the dynamics until convergence. Then we measure the overlap between the final state and the memory pattern. We keep increasing the number of flipped units until the final state has an overlap smaller than ≥ 0.85 with the memory state. We define the normalized basin size as the maximal number of flips allowing for a large overlap divided by 2*f N*, the maximal number of flips. Results are shown in the SI.

### C. Noisy synaptic dynamics

The synaptic dynamics in the presence of Gaussian noise is presented in equation (13). It is straightforward to show that a solution to the equation can be written as:

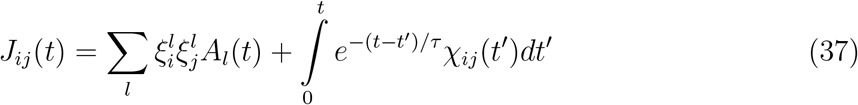

with *A*_*l*_(*t*) obeying eq. (7) as before. The nonlinear effect of the noise arises through the self consistent requirement, that the rehearsal rate of memory *l* is proportional to the basin of attraction size of this memory, which depends on *A*_*l*_ and on Δ. We calculate Δ with the injected noise (eq. (14)) the same way we calculated Δ without noise above. Here there is a non trivial mixed term involving the average *A*_*l*_(*t*)*χ*_*ij*_(*t*), which we found numerically to be negligible for the parameter range we are interested in.

### D. Synaptic dilution

The random silencing is done by multiplying the connectivity matrix *J*_*ij*_ by a binary matrix:

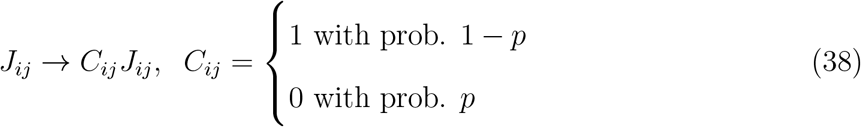

We would like to calculate the effect of the dilution on the memory efficacies dynamics, and for that we need to find the effect on *A*_*l*_(*t*) and on Δ(*t*). Let us calculate the local field near memory *l* as before:

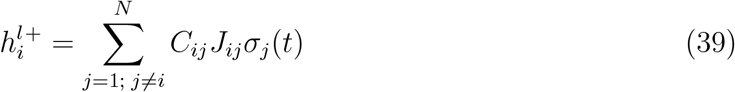

Taking the mean over memories and over *C*_*ij*_ realizations (denoted by []) we get:

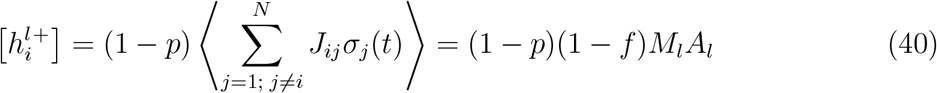

Here ⟨ ⟩ denotes average over memories realizations. As one can see, the efficacies are scaled by a factor of 1 − *p*. We assumed here we can neglect correlations between *A*_*l*_ and *C*_*ij*_. The second moment:

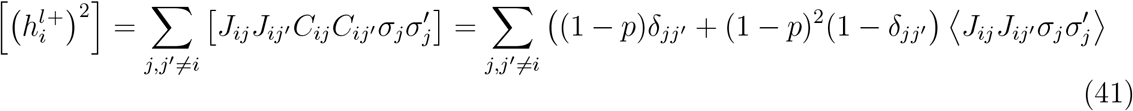

There are two contributions arising from the *C*_*ij*_ randomness. Now, when calculating the local field variance, the term proportional to (1 − *p*)^2^ is exactly canceled by the squared mean, and we are left with:

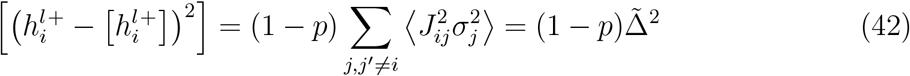

where 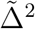 is the local field variance without dilution.

Now, we obtain the efficacies modified dynamics by using these expressions for the signal and noise to calculate the basins of attraction sizes as before.

### E. Non-uniform characteristic decay time

Assuming synapse *J*_*ij*_ has a decay rate *ϵ*_*ij*_, and all memories have unit initial efficacy. Memory *l* appears for the first time at time *l*. The learning dynamics is:

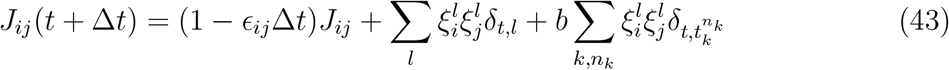

In continuous time,

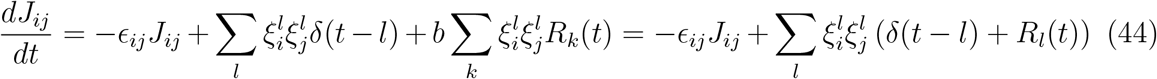

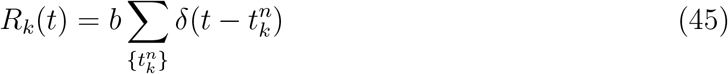

Assuming all synapses starts at zero value, the solution can be written as:

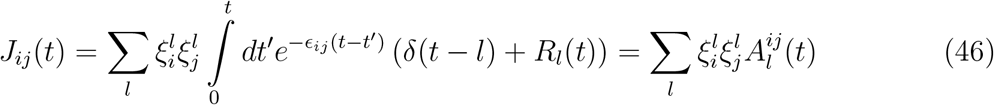

Let us define efficacies

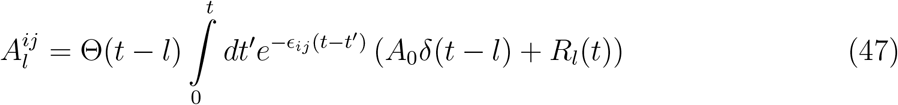

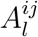 obeys the differential equation:

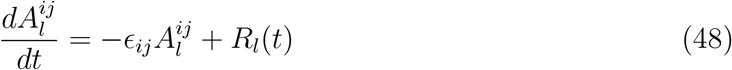

With 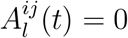 for *t < l* and 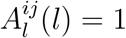.

Given that the decay rates have a probability density *ρ*(*ϵ*), let us define:

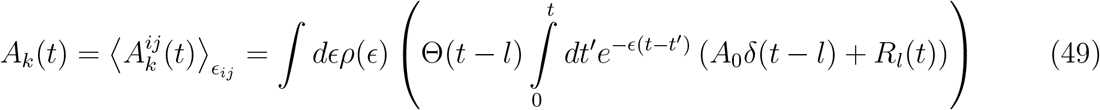

and

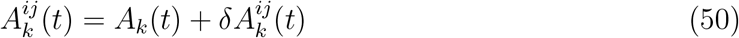

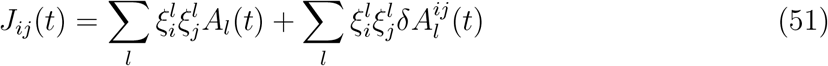

Including normalization and sparseness considerations,

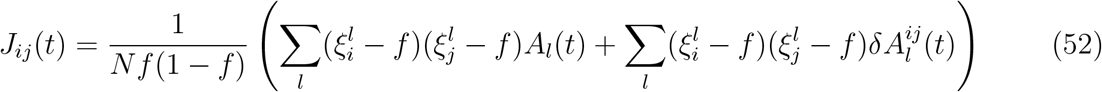

Now let us calculate the mean local field on neuron *i* in a state near memory state *k*, and assume 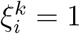:

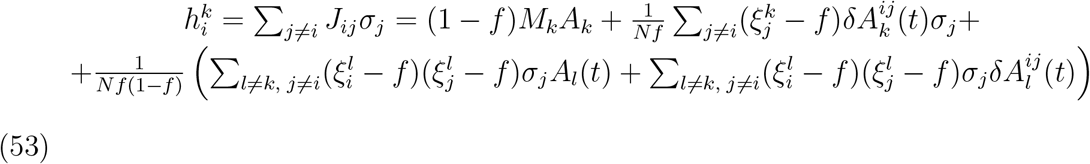

Taking an average over the memories realizations and the decay rates, we get:

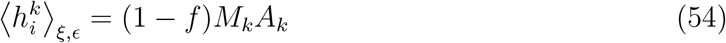

And the variance:

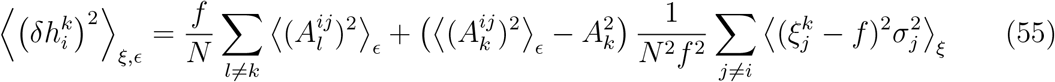

The second term does not include summation over all memories, and therefore it is negligible for large N values (the first term is *O*(1) while the second is *O*(*N* ^−2^). This leads to eq.(19).

#### a. Power law τ distribution

For each synapse we generated synaptic decay characteristic times from a power law (Pareto) distribution with density:

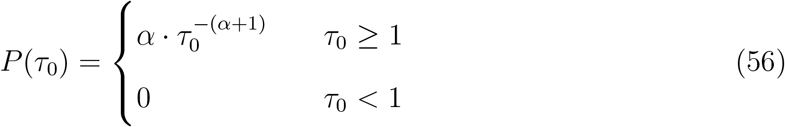

In this distribution, for *α <* 1 the mean diverges. We scaled the resulting 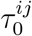 values by a uniform factor: 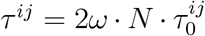. We fixed *ω* value for the average number of rehearsals per mean decay time *R*_0_, and used it to set the *λ* parameter by dividing *R*_0_ by the empirical average of the generated decay times. Next we solved the stochastic differential equations (48) numerically. The rehearsals are generated with time dependent rates proportional to the basin of attraction size, now as a function of the average and variance of the memory efficacies over all synaptic timescales.

