## Supplementary information for "Stochastic Consolidation of Lifelong Memory"

### Stochastic Consolidation of Lifelong Memory - Supplementary Information

**CONTENTS**


I. Basins of attraction 3

A. numerical calculation 3

B. Approximated equation for $A_c(f, \Delta)$ 5

C. Fixed threshold 6

II. Efficacies fluctuations around the fixed point $A_{fp}$ 11

III. The effective forgetting time, consolidation probability and and capacity 12

IV. Synaptic dynamics perturbations - additional results 17

V. Transient synaptic dilution 20

VI. Deterministic model 21

References 24

### I. BASINS OF ATTRACTION

#### A. numerical calculation

We use a mean field theory to approximate the basin of attraction of a memory pattern as a function of the memory efficacy $A_l(t)$ and of the global interference noise amplitude $\Delta(t) = f/N \sum_k A_k^2(t)$ (Fig. S1, see *methods* in the main text). To check the validity of our approximation, we ran full network simulations, where random activation patterns (dimension $N$, sparseness $f$) were generated and stored in the network via the synaptic dynamics (Eq. (1) in the main text), where we used the basin of attraction approximated by the mean field calculation. Then, for a given memory, we generated a perturbed initial condition in the following way: $k$ active units ($0 \leq k \leq fN$) were chosen randomly and flipped to inactive state, and $k$ inactive units in the memory state were chosen randomly and flipped to active state (so that the total activation of $fN$ is maintained). Then, we initialized the network dynamics at that state, ran it until convergence, and measured the overlap between the final state and the original memory state. We kept increasing $k$ until the final overlap dropped beneath 0.85 (or $k$ reached $fN$). We repeated this process 50 times, in each we chose randomly the units to be flipped. We define the basin of attraction size as $k_{max}/fN$, where $k_{max}$ is the average (over trials) of the maximal $k$ for achieving final overlap of 0.85. In Fig. S2 we plot the basin of attraction as a function of the efficacy $A$ for both the full simulation calculation and for the mean field approximation for several values of $N$. As one can see, the prediction is good, and especially the prediction of the critical efficacy, where the basin size vanishes. As shown in Fig. (3) in the main text, the forgetting curves obtained from the full network simulations and the mean field approximation fit perfectly, regardless of the small deviations seen in the basins size.

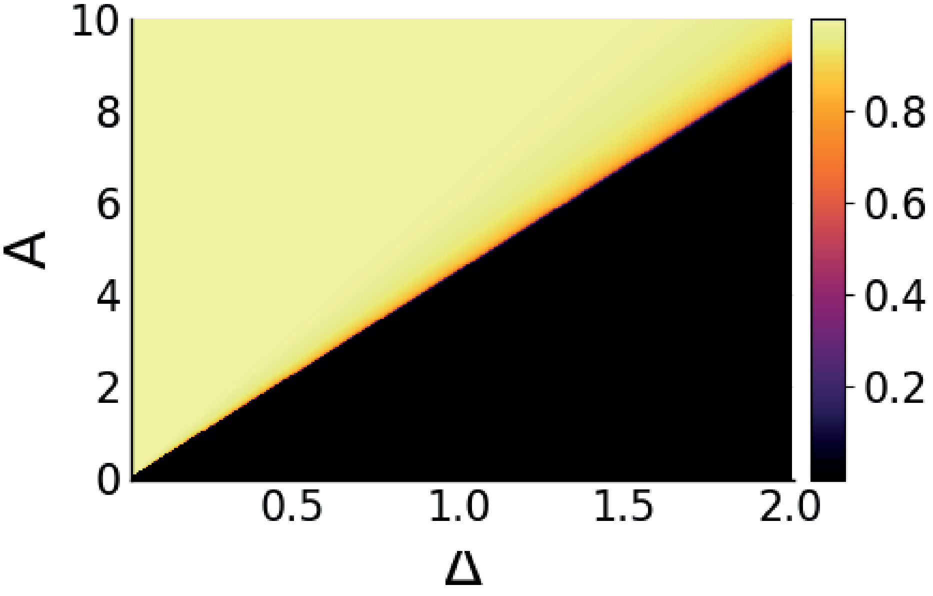


FIG. S1. Basin of attraction size (color code) as a function of the memory efficacy $A$ and the interference noise $\Delta$. Here $f = 0.01$.

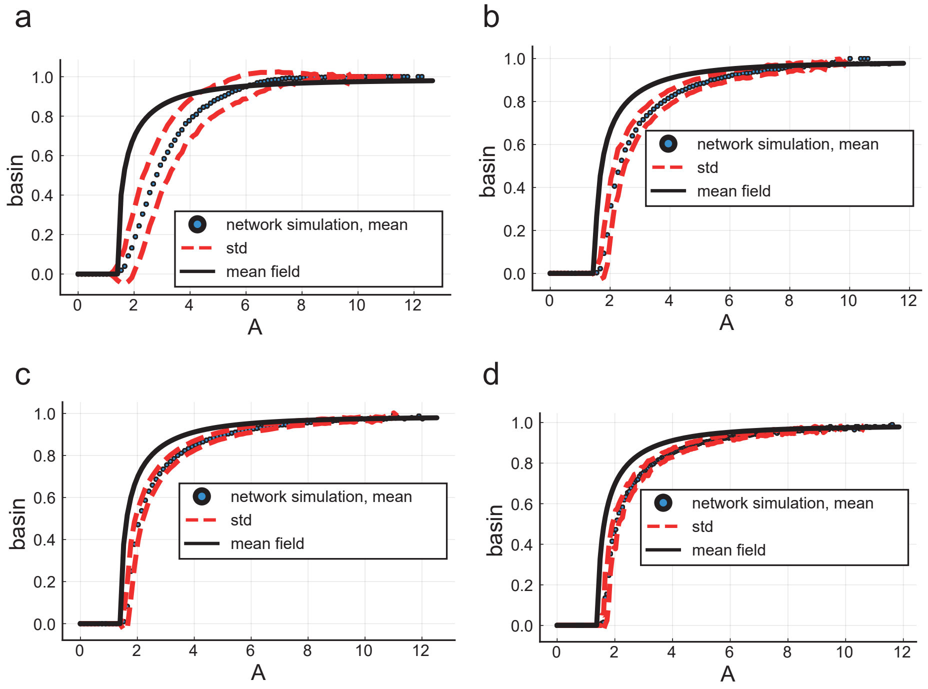


FIG. S2. Basin of attraction VS. memory efficacy. Basin size calculated using the mean field theory (black) and full numerical simulations (blue points, with std in red dashed lines), for different values of $N$: $N = 1000,\ 4000,\ 8000,\ 10000$ for **a, b, c, d** respectively.

##### B. Approximated equation for $A_c(f, \Delta)$

Assuming $f \ll t$ and large overlap ($M \approx 1$, and therefore $f_- \ll 1,\ 1-f_+ \ll 1$), equations (31) and (32) in the main text can be approximated by:

$$1 - f_+ \approx H\left(\frac{A_l - \theta}{\Delta}\right) \approx \frac{\Delta}{A_l - \theta}\exp\left(-\frac{(A_l-\theta)^2}{2\Delta^2}\right) \to -\log(1 - f_+) \approx \frac{(A_l-\theta)^2}{2\Delta^2} \quad \text{(S1)}$$

$$f_- \approx \frac{\sqrt{\Delta}}{\theta}\exp\left(-\frac{\theta^2}{2\Delta^2}\right) \to -\log(f_-) \approx \frac{\theta^2}{2\Delta^2} \quad \text{(S2)}$$

Now, using the fixed activation assumption:

$$f = \frac{1}{N}\sum_{i=1}^{N}\sigma_i(t) = ff_+ + (1-f)f_- \quad \text{(S3)}$$

$$\frac{f_-}{1-f_+} = \frac{f}{1-f} \quad \text{(S4)}$$

$$-\log(1-f_+) = \frac{A_l^2 - 2\theta A_l + \theta^2}{2\Delta^2} = \frac{A_l^2 - 2\theta A_l}{2\Delta^2} - \log(f_-) \quad \text{(S5)}$$

$$\log\left(\frac{f_-}{1-f_+}\right) = \frac{A_l^2 - 2\theta A_l}{2\Delta^2} \quad \text{(S6)}$$

$$\log\left(\frac{f}{1-f}\right) \approx \log(f) = \frac{A_l^2 - 2\theta A_l}{2\Delta^2} \quad \text{(S7)}$$

we obtain an expression for the threshold $\theta$:

$$\theta = \frac{A}{2} + \frac{\Delta^2}{A}\log(1/f) \quad \text{(S8)}$$

Demanding that the efficacy is larger than $\theta$, we obtain:

$$A_c \approx \sqrt{2\log(1/f)}\Delta \quad \text{(S9)}$$

and therefore, $a(f) = \sqrt{2\log(1/f)}$.

By solving numerically the full fixed point equation in the main text (36) without taking $M \to 1$ , we found that the approximated expression gives the right functional behavior, but required a constant numerical correction (Fig. S3):

$$A_c = 1.44\sqrt{2\log(1.9/f)}\Delta \quad \text{(S10)}$$

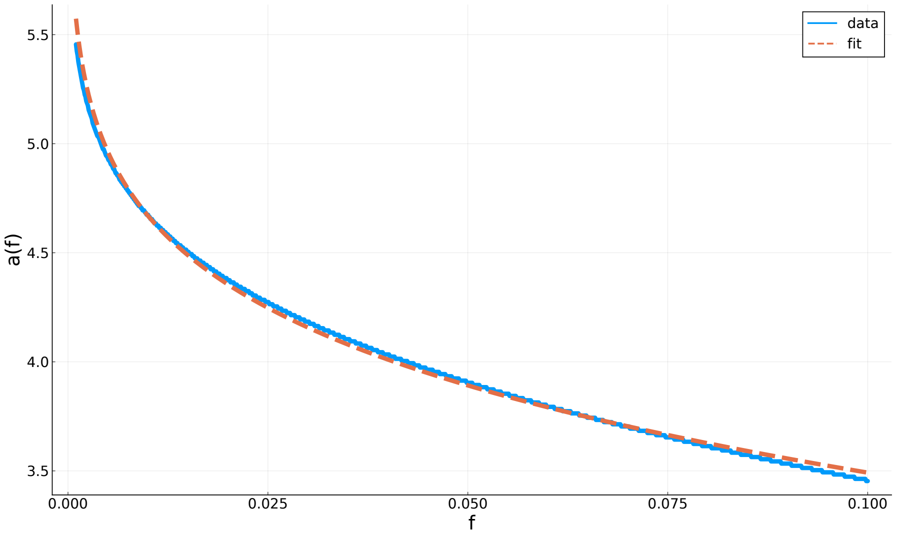


FIG. S3. $a(f)$ as calculated numerically (solid line) and the analytical approximation (dashed line) as a function of $f$.

##### C. Fixed threshold

When assuming fixed threshold instead of fixed activation level, the basin calculation is modified: instead of equation (25) in the main text we have an equation for the mean activation level:

$$\mu(t) = \frac{1}{N}\sum_{i=1}^{N}\sigma_i(t) = f f_+(t) + (1-f) f_-(t) \tag{S11}$$

By a similar calculation as in the *Methods* section, using equations (31), (32) and (27) in the main text together with eq. (S11) we can write the overlap and activation dynamics as:

$$\begin{aligned} M_l(t+1) &= G_1(M_l(t), \mu(t); A_l, \Delta, \theta) \\ \mu(t+1) &= G_2(M_l(t), \mu(t); A_l, \Delta, \theta) \end{aligned} \tag{S12}$$

where

$$G_1(M_l, \mu; A_l, \Delta, \theta) = H\left(\frac{\theta - A_l(1-f)M_l}{\sqrt{\mu}\Delta}\right) - H\left(\frac{\theta + A_l M_l f}{\sqrt{\mu}\Delta}\right) \tag{S13}$$

and

$$G_2(M_l, \mu; A_l, \Delta, \theta) = f \cdot M_l(t) + H\left(\frac{\theta + A_l M_l f}{\sqrt{\mu}\Delta}\right) \tag{S14}$$

For a given threshold $\theta$ the basin of attraction size is a function of two variables, memory efficacy $A_l$ and the interference noise $\Delta^2 = \frac{1}{N}\sum_l A_l^2$. We estimate the basin size for given

values of $A_l$, $\Delta$ and $\theta$ in a similar way to the fixed activation scenario (*Methods*): First, we run the dynamics described by equation (S12) from initial condition $M(0) = 1,\ \mu(0) = f$ until convergence. If $M$ at convergence ($\equiv M_f$) $\leq 0.85$, the basin size is zero. Else, we repeat the process with smaller $M(0)$ value until either $M_f$ is smaller than 0.85 or $M(0)$ is smaller than 0.01. We define the basin size as the distance between the unperturbed overlap (starting at $M(0) = 1$) and the minimal $M(0)$ for which $M_f > 0.85$ (or zero if we reach $M(0) < 0.01$). Results are shown in Fig. S4. The main qualitative differences are that $A_c$ does not go to zero for $\Delta = 0$ , because even in the absence of interference, the fixed threshold puts a lower bound on the value of the local fields that will allow activation of neurons. As a result, $A_c$ is finite at low $\Delta$ and increases very slowly with it. However, above a critical value of $\Delta$ fluctuations are strong enough to allow neurons that are off in the memory state to turn on, effectively making all memories unstable (similar to the catastrophic forgetting in the classical sparse Hopfield model [**?** ]). The system's capacity depends now on the value of the threshold (relative to the encoding strength $A(0) = 1$). On one hand, increasing the threshold size increases the minimal efficacy required for activation, hence $A_c$ increases; on the other hand it increases the critical $\Delta$ value at which all memories become unstable. Therefore we should expect an optimal threshold value for maximizing the capacity [**?** ]. Indeed, for a given $b,\ \lambda\tau$ we calculate the capacity for different threshold values and show that there is an optimal value (Fig. S6b), where capacity is similar to the adaptive threshold (constant activation) setting.
Memory efficacies dynamics and forgetting curve shapes are qualitatively similar to the constant activation setting (Fig. S6a).

When modeling memory deficits due to perturbations, we used the fixed threshold scenario in order to account for potential memory loss due to disruption of the level of activity in the network; specifically, allowing for memory loss due to insufficient activation (in the case of synaptic dilution) or due to over-activation (in the case of noisy synaptic dynamics). The fixed threshold was chosen as the one that maximizes the capacity in the control case (before perturbations). For the chosen parameters $\tau/(2N) = 0.01,\ \lambda\tau = 10,\ b = 0.2$, the optimal threshold is $\theta = 0.36$ (Fig. S6b). In the threshold adaptation paragraph we adapted the threshold to a value maximizing capacity in the presence of the perturbation.

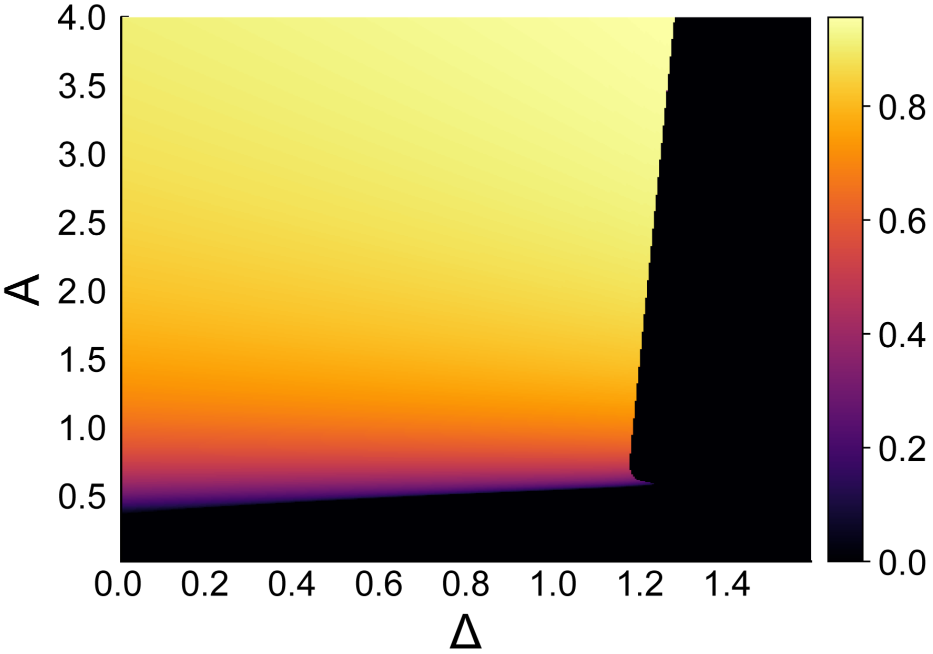


FIG. S4. Basin of attraction size (color code) as a function of the memory efficacy $A$ and the interference noise $\Delta$. Here $\theta = 0.36,\ f = 0.01$.

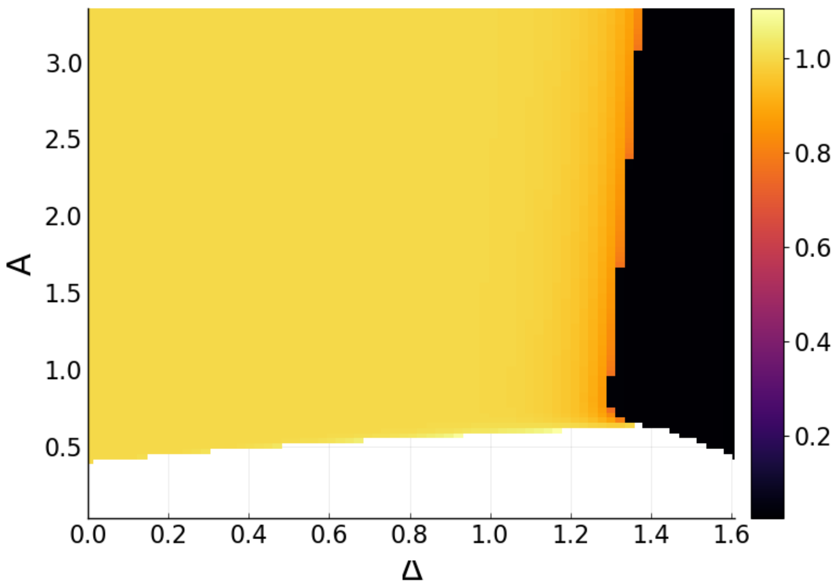


FIG. S5. Normalized population activation (color code) as a function of the memory efficacy $A$ and the interference noise $\Delta$. The normalized activation is $f$ (the activation level of the memory patterns) divided by the activation $(\mu)$ (white - low activation compared to the memory patterns. black: high activation). Here $\theta = 0.36,\ f = 0.01$.

a

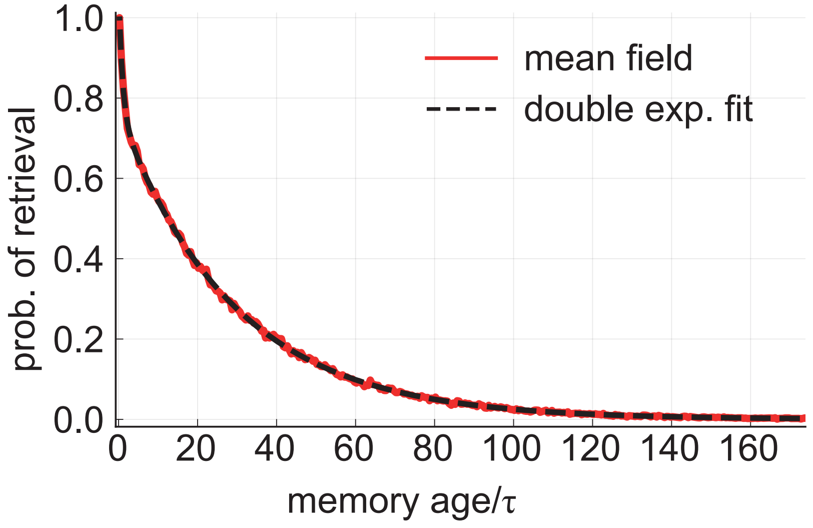


b

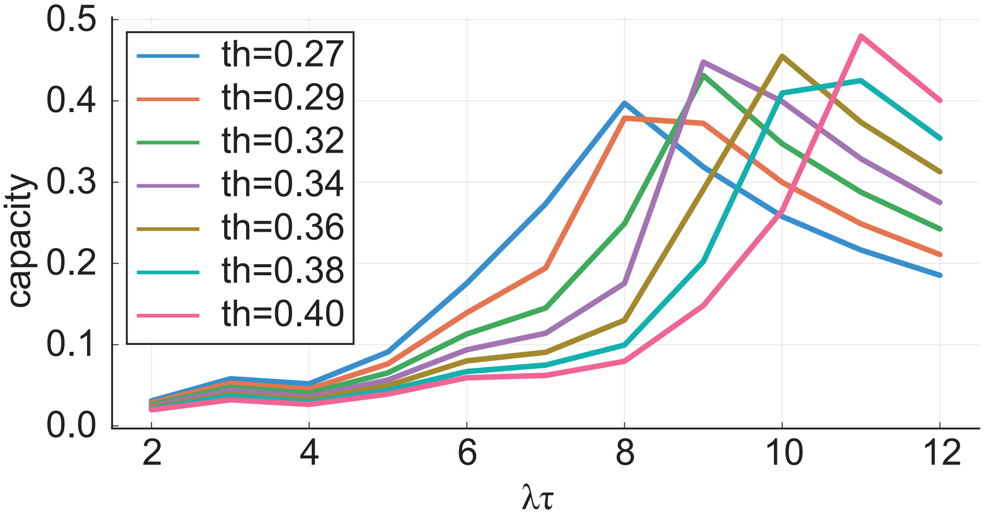


FIG. S6. **a** Retrieval probability Vs. memory age. Mean field calculation results for $N = 1000$, $\lambda\tau = 10$, $b = 0.2$, $\theta = 0.36$. Fit to a double exponential is shown, the long time scale: $\tau_c \approx 30\tau$. **b** capacity Vs. $\lambda\tau$. $N = 1000$, $\lambda\tau = 10$, $b = 0.2$, $\theta = 0.36$, $b = 0.2$.

#### II. EFFICACIES FLUCTUATIONS AROUND THE FIXED POINT $A_{fp}$

At the steady state (after $\Delta$ had saturated), after a memory gets consolidated, its efficacy fluctuates around the fixed point $A_{fp}$ until a large enough fluctuation drives it beneath $A_c$, and then it decays exponentially to zero. The stochastic rehearsals driving the random walk around $A_{fp}$ can be approximated by a Poisson process with rate:

$$\tilde{\lambda} = \lambda F(A_{fp}/\Delta) \tag{S15}$$

where $F(A_{fp}/\Delta)$ is the basin of attraction size function evaluated at $A_{fp}$ and the steady state $\Delta$. Now, $n$ rehearsal events after consolidation, the value of the efficacy can be written as:

$$A(t) = b\sum_{i=1}^{n} \exp((t_i - t)/\tau) \tag{S16}$$

where $t_i$ is the time of rehearsal event $i$. Let us define $\Delta t_i \equiv t_i - t$ and the total time since consolidation

$$T \equiv \sum_{i=1}^{n} \Delta t_i \approx \frac{n}{\tilde{\lambda}} \tag{S17}$$

Now, define

$$y_i = \exp(\Delta_i/\tau),\ 0 < y_i \leq 1 \tag{S18}$$

which are independent random variables ($\Delta t_i$ are independent) , and $A(t) = b\sum_i y_i$. The probability density of $y_i$:

$$p(y) = p(\Delta t)\frac{d\Delta t}{dy} = \frac{\tau}{T}\frac{1}{y} = \frac{\tilde{\lambda}\tau}{n}\frac{1}{y} \tag{S19}$$

We used the fact that $\Delta t_i$ are uniformly distributed on $T$. Now, using the condition for a fixed point,

$$\tau\tilde{\lambda} = \tau\lambda F(A_{fp}/\Delta) = A_{fp}/b \tag{S20}$$

so that

$$p(y) = \frac{A_{fp}}{bn}\frac{1}{y} \tag{S21}$$

Now we can calculate the mean and variance of $A(t)$

$$\langle A(t)\rangle = bn \cdot \langle y\rangle = A_{fp} \tag{S22}$$

$$\mathrm{var}(A(t)) = b^2 n \cdot \mathrm{var}(y) = \frac{bA_{fp}}{2} - \frac{A_{fp}^2}{n} \to \frac{bA_{fp}}{2} \tag{S23}$$

The last limit assumes large number of rehearsals $n$.

#### III. THE EFFECTIVE FORGETTING TIME, CONSOLIDATION PROBABILITY AND AND CAPACITY

After $A_c$ saturates, the behavior of every memory efficacy is a random walk with an attraction to $A_{fp}$. This process will continue until the first time the efficacy will cross $A_c$, and from that moment the memory is forgotten, and will decay exponentially. The forgetting curve is essentially the distribution of first passage times for this stochastic process. For Gaussian noise it is known [1, 2] to be an approximate exponential distribution, with mean first passage (forgetting) time:

$$\tau_{\mathrm{c}} \approx \tau \mathrm{exp}\left(\frac{(A_{fp}-A_c)^2}{2\sigma^2}\right) \tag{S24}$$

Where $\sigma^2$ is the variance of the driving Gaussian process. In our case the efficacy dynamics is driven by non-homogeneous Poisson noise, and there is no closed form solution for the mean first passage time from $A_{fp}$ to $A_c$ we are aware of. To get an approximated and insightful expression for the mean life time of consolidated memories, we work as follows: First, in the absence of rehearsals, a memory will drop from $A_{fp}$ to $A_c$ over $\tau \log A_{fp}/A_c$ time. In a homogeneous Poisson process with rate $\lambda$, the chance of not having events for that time is $\exp\left(-\lambda\tau\log\left(A_{fp}/A_c\right)\right) = (A_c/A_{fp})^{\lambda\tau}$. However, in our model the rehearsal rate is not homogeneous. Therefore, we estimate the initial point and the rehearsal rate at an intermediate point, which we find empirically to give a good fit to the data. We then approximate the mean forgetting time as $\tau$ times the inverse of the probability function:

$$\tau_c \approx \tau X^{\lambda\tau F(X)},\ X \equiv \frac{0.25b\lambda\tau + 0.75A_c}{A_c} \tag{S25}$$

We found empirically that the first passage time distribution is again exponential (Fig. 3a,b in the main text). Given the exponential shape of the forgetting curve and $\tau_{\mathrm{c}}$, we can evaluate the capacity $n_r$, the number of memories which are retrievable at the steady state, as an integral over the forgetting curve (which equals $\tau_{\mathrm{c}}$), multiplied by the chance of a memory to get consolidated (or, equivalently, reach $A_{fp}$ at some time point):

$$n_r \simeq p_c \tau_{\mathrm{c}} \tag{S26}$$

where $p_c$ is the probability of a memory efficacy to get consolidated and join the efficacies fluctuating around $A_{fp}$.

We approximate $p_c$ by assuming that memories that don't consolidate are the ones free-falling exponentially from the initial point $A_0$ to the critical point $A_c$. The time it takes is $\tau\log(A_0/A_c)$, and therefore, due to the Poisson statistics (we again use an intermediate point, due to the non homogeneous rate),

$$p_c(A_0, A_c) \approx 1-\exp\left(-\lambda F(A_0/A_c)\tau\log(0.5(A_0 + A_c)/A_c)\right) = 1-(A_c/0.5(A_0+A_c))^{\lambda\tau F(0.5(A_0+A_c)/A_c)} \tag{S27}$$

Here $F(x)$ is the basin of attraction size.

As we have seen above the long life time of consolidated memories as well as the memory capacity depend on the steady state critical efficacy $A_c$ and the fixed point efficacy $A_{\mathrm{fp}}$. In fact, knowing $A_c$, the fixed point efficacy can be determined by the fixed point equation $A_{\mathrm{fp}} = b\lambda\tau F(A_{\mathrm{fp}}/A_c)$, and is in general close to $b\lambda\tau$. Then, $\tau_{\mathrm{c}}$ (and therefore the capacity) can be determined by eq. (S25). To evaluate $A_c$ or equivalently the interference noise at steady state, we evaluate the second moment of all efficacies via the stationary efficacy histogram, Fig. 2c. One contribution is from the central 'bump' corresponding to consolidated memories fluctuating around the fixed point. These efficacies contribute approximately $n_r(A_{\mathrm{fp}}^2+0.5A_{\mathrm{fp}})$ where the second term is the contribution of the Poisson variance around the mean. Another contribution is from memories with efficacies smaller than $A_c$ which decay to zero, hence contribute $0.5A_c^2\tau$. Finally, when $1 \gg A_{\mathrm{fp}}$ there is a long tail outside the center peak, coming from the initial decay of the high efficacies toward the fixed point, contributing roughly $0.5\tau$ (when $A_{\mathrm{fp}}$ is not very small compared to unity, this term is negligible compared to the rest). Thus, we approximate $A_c$ as

$$A_c^2 = \frac{a^2(f)f}{2N}\left[p_c\tau_{\mathrm{c}}(2A_{\mathrm{fp}}^2 + A_{\mathrm{fp}}) + \tau(A_c^2+1)\right] \tag{S28}$$

Together with equations (S25), and (S27), we have a set of self consistent equations for the macroscopic variables of the theory. When consolidated memories provide the dominant contribution (which is the regime we are interested in, where the situation is qualitatively different than pure forgetting), the consolidation probability is close to unity and $A_{fp} \approx b\lambda\tau$, $A_c$ can be approximated as:

$$A_c^2 = \frac{a^2(f)f}{N}\tau_{\mathrm{c}}(b\lambda\tau)^2 \tag{S29}$$

Under these assumptions, $\tau_c \approx n_r$ can be approximated as:

$$\tau_c \approx n_r \propto \tau\left(\frac{b\lambda\tau}{A_c}\right)^{\lambda\tau} \tag{S30}$$

Now, using eq. (S29), we get:

$$\tau_c \approx \tau \left( \sqrt{\frac{N}{a^2(f) f \tau_c}} \right)^{\lambda\tau} \rightarrow \tau_c \approx \tau^{\frac{1}{1+0.5\lambda\tau}} \left( \frac{N}{a^2(f) f} \right)^{\frac{\lambda\tau}{\lambda\tau+2}} \tag{S31}$$

The theory gives a good fit to simulation results (Fig. and Fig.4 in the main text) and allows us to observe the relations between model parameters ($\tau$, $\lambda, b$, $f$, $N$) and the system's behavior.

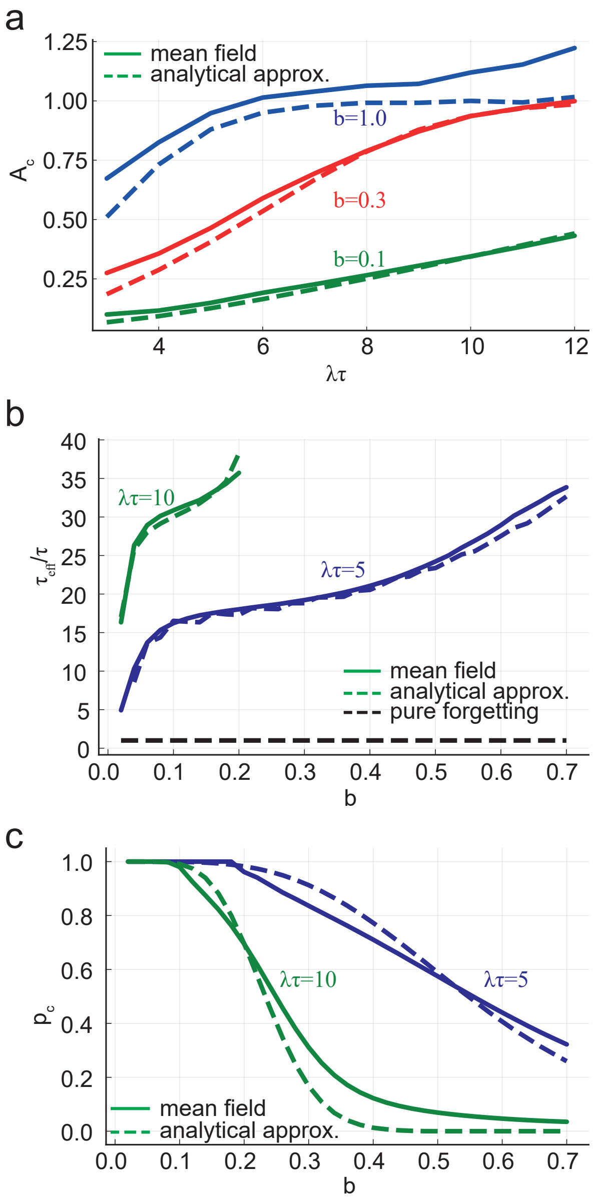


FIG. S7. *Analytical theory vs simulation results.* **a** $A_c$ vs. $\lambda\tau$ for different $b$ values. **b** $\tau_c/\tau$ vs. $b$ for different $\lambda\tau$ values. **c** Consolidation probability vs. $\lambda\tau$ for different $b$ values. Other parameter values: $N = 8000,\ f = 0.01,\ \tau = 160$

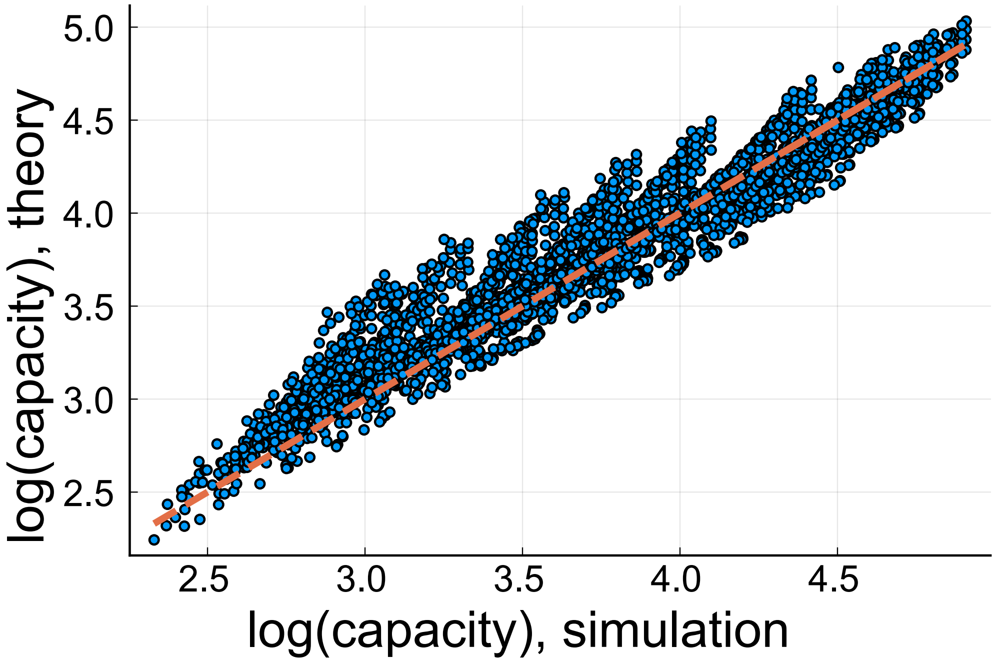


FIG. S8. $\tau_c$ from numerical simulations vs. the analytical approximation (blue points). The $x = y$ line is shown. Here $N = 8000,\ f = 0.01,\ \tau = 160$. Results are shown for $\lambda\tau$ values between 4 and 12 and $A_0$ values between 2 and 40.

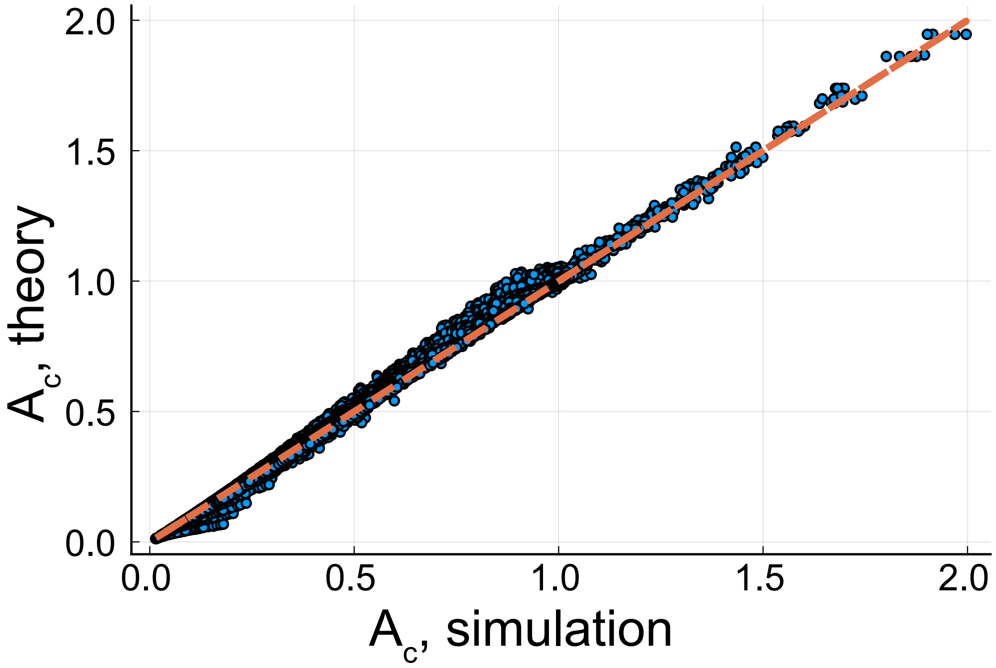


FIG. S9. $A_c$ from numerical simulations vs. the analytical approximation (blue points). The $x = y$ line is shown (dashed line). Here $N = 8000,\ f = 0.01,\ \tau = 160$. Results are shown for $\lambda\tau$ values between 3 and 12 and $b$ values between 0.1 and 5.

#### IV. SYNAPTIC DYNAMICS PERTURBATIONS - ADDITIONAL RESULTS

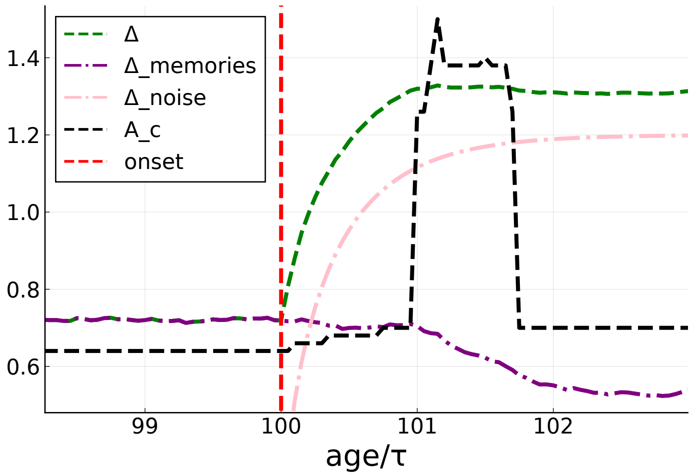


FIG. S10. *$A_c$ and $\Delta$ dynamics in the presence of additive noise*: Green dashed line: total $\Delta$. Black dashed line: $A_c$. Purple dash-dot line: Memories' efficacies contribution to $\Delta$. Pink dash-dot line: noise contribution to $\Delta$. After the noise onset, $\Delta$ rises due to the additive noise contribution until a critical $\Delta$ is reached, where $A_c$ rises abruptly. Then, due to lack of rehearsals, memory efficacies decay and $\Delta$ is reduced, until $A_c$ settles to a new value (a bit higher than the original value) that allows for rehearsals.

Parameters: $N = 8000,\ \tau = 160,\ \lambda\tau = 10,\ b = 0.2$

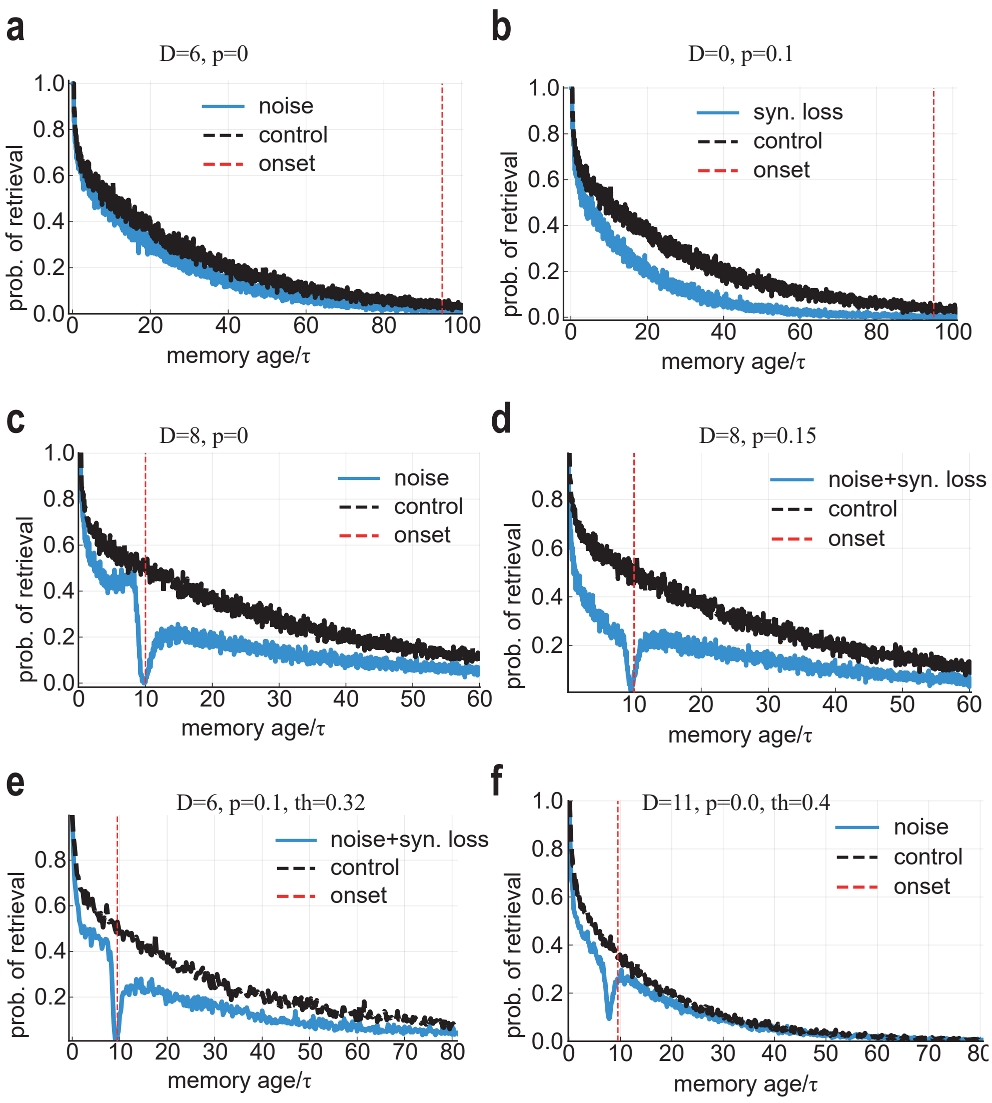


FIG. S11. Perturbations and memory deficits - additional results. **a** Retrieval probability vs. memory age with noisy synaptic dynamics ($D = 6$), at time $95\tau$ after the noise onset. The control (black) is simulated with noiseless dynamics. **b** Retrieval probability vs. memory age for random synaptic dilution ($p = 0.1$), at time $95\tau$ after the noise onset. **c** Retrieval probability vs. memory age with noisy synaptic dynamics ($D = 8$) and no dilution, at time $10\tau$ after the noise onset. **d** Combination of synaptic dilution and noisy synaptic dynamics, $D = 8$ and $p = 0.15$.. **e** Low threshold compared to optimal: here $\theta = 0.32$, and $D = 6$, $p = 0.1$. **f** High threshold compared to optimal: here $\theta = 0.4$, and $D = 11$, no dilution.

Parameters: $N = 8000,\ \tau = 160,\ \lambda\tau = 10,\ b = 0.2$

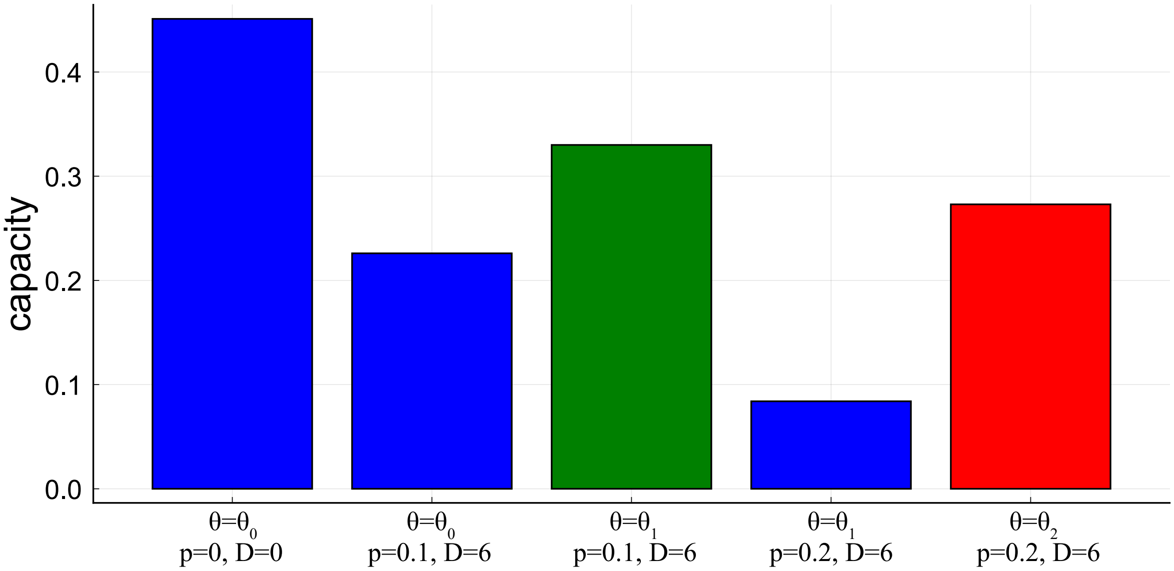


FIG. S12. Effect of threshold adaptation with synaptic dilution and additive noise. Blue bars are for threshold optimized for the noiseless case ($\theta_0 = 0.36$). red bar is for threshold optimized for low dilution (p=0.1) and synaptic noise ($D = 6$) ($\theta_0 = 0.33$). green bars are for threshold optimized for high dilution (p=0.2) and synaptic noise ($D = 6$) ($\theta_0 = 0.31$). Parameters: $N = 8000,\ \tau = 160,\ \lambda\tau = 10,\ b = 0.2$

#### V. TRANSIENT SYNAPTIC DILUTION

In the main paper we present results for ongoing of a fraction of the synapses. Here, we show what happens if there is a transient event of silencing (for example, a temporal blockage of a blood vessel, etc). In Fig.S13 we show the forgetting curve for a strong transient event lasting for $2\tau$ time. There is a significant reduction in retrieval of memories entered just before the onset (temporally graded retrograde amnesia) and a tremendous reduction of retrieval probability for memories learned during the event. This is similar to the phenomenology of *Transient Global Amnesia* (TGA), where patients suffer from a whole day during which they have hard time retrieve past memories, and after the episode is over there is a strong amnesia of events that happened during the episode, and of events that happened just before the event [3].

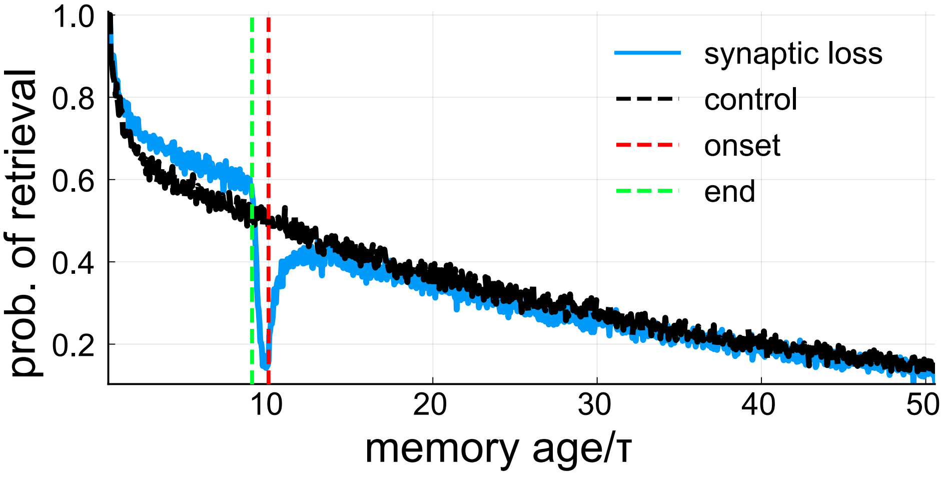


FIG. S13. Retrieval probability vs memory age for transient synaptic silencing (blue). 30% of the synapses were silent for $2\tau$ time, and then recovered. Other parameters as in Fig.6d in the main text.

#### VI. DETERMINISTIC MODEL

When replacing the Poisson rehearsals process by its average, a deterministic model for the memory efficacies arises (In this part we will use $b = 1$, and vary $A_0$):

$$\frac{dA_l}{dt} = -\frac{1}{\tau}A_l + \lambda F(A_l/\Delta), \quad A_l(t = l) = A_0, \quad A_l(t < l) = 0 \tag{S32}$$

$$\Delta^2(t) = \frac{f}{N}\sum_l A_l^2(t) \tag{S33}$$

Define $A_c(t) = \alpha(f)\Delta(t)$ such that

$$F(\frac{A \leq A_c}{\Delta}) = 0 \tag{S34}$$

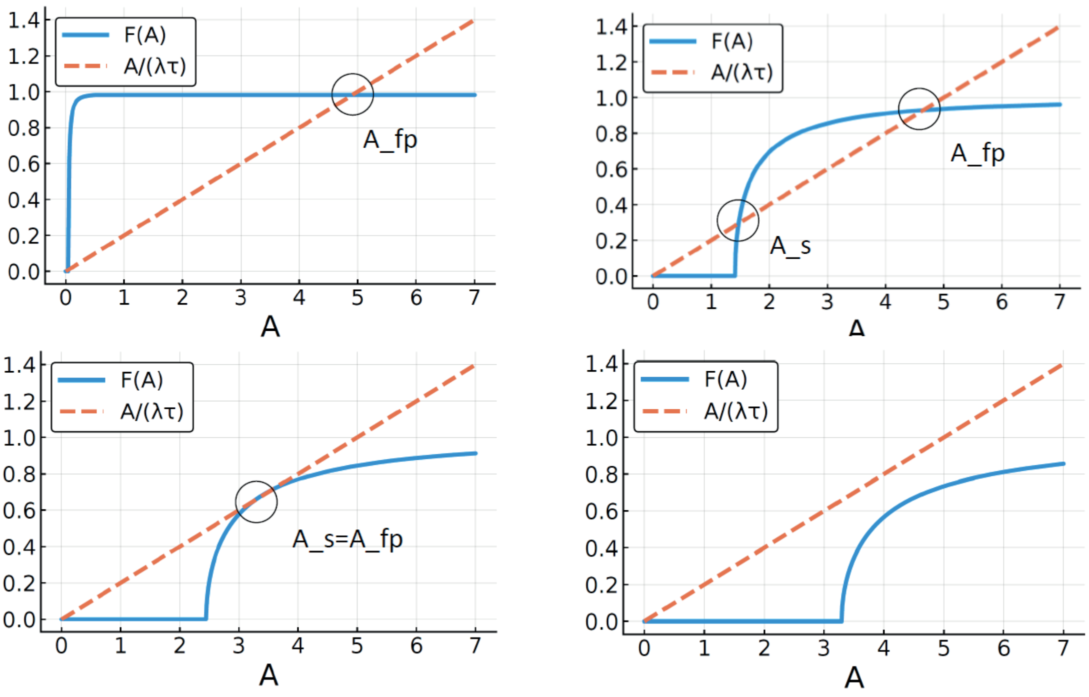


FIG. S14. $F(A/\Delta)$(blue) and $A/(\lambda\tau)$ (dashed red) for different values of $\Delta$ (close to zero in the top left, intermediate in the top left, the critical $\Delta$ in the bottom left and large $\Delta$ in the bottom right. The nonzero fixed points are marked.

Initially all A's are zero, and so is $\Delta$. At that phase, every A that enters grows towards $A_{fp}$ (which is initially $\lambda\tau$, and decrease slowly when $\Delta$ increase). While A's accumulate at $A_{fp}$, $\Delta$ grows.

What will happen next depends on $A_0$: for small enough $A_0$, $\Delta$ will grow until $A_s(\Delta) = A_0$. From that point on, all the new A's will decay exponentially, and all the past A's will stay at $A_{fp}$, determined by the $\Delta$ at that point (Fig. S15 top). We can write an equation relating $\Delta$ to the number of A's that will stay in $A_{fp}$: by:

$$\Delta^2 \approx n\frac{f}{N}A_{fp}^2 + \frac{f}{2N}A_0^2\tau \tag{S35}$$

The first contribution comes from the memories accumulated at $A_{fp}$ (n is their number) and the second contribution comes from the decaying memories.

For large enough $A_0$, $\Delta$ will grow until $A_s(\Delta) = A_{fp}(\Delta) < A_0$. When that happens, $A_{fp}$ looses its stability, and all the A's accumulated there will start decaying exponentially. That will cause $\Delta$ to decrease, restoring the stability of $A_{fp}$, and memories will start accumulating there again, and $\Delta$ will start increasing again - only until $A_s(\Delta) = A_{fp}(\Delta)$. This process will continue periodically forever, constantly storing new groups of memories at $A_{fp}$ and deleting them, replacing them with a new group (Fig.S15, bottom).

The value of the maximal $\Delta$ will be determined by $A_s(\Delta) = A_{fp}(\Delta)$, or:

$$\frac{1}{\lambda\tau}A = F(A/\Delta)$$

$$\frac{1}{\lambda\tau} = \frac{1}{\Delta}\frac{d}{dA}F(A/\Delta)$$

Solving these equations gives also the critical $A$ value, where the bifurcation from fixed point behavior to a limit cycle oscillatory behavior as a function of $A_0$ happens.

In the stochastic case studied in the main paper, a noisy version of the fixed point phase is observed for small enough $A_0$ values. However, in the stochastic case the limit cycle phase doesn't exist, due to the large fluctuations around $A_{fp}$. Nevertheless, if we replace the Poisson noise by a Gaussian noise with small enough amplitude, a noisy limit cycle behavior can be observed (not shown).

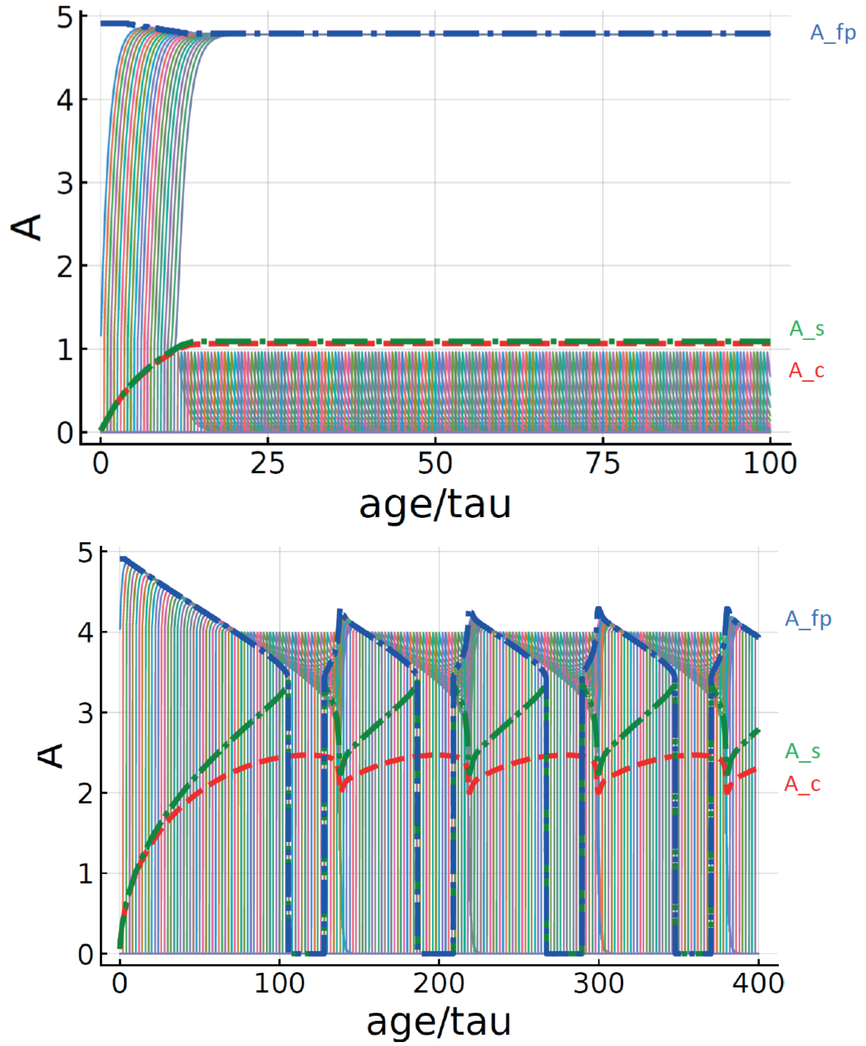


FIG. S15. Efficacies trajectories vs. time for the deterministic model with small $A_0$ (top) and large $A_0$ (bottom).

[1] H. A. Kramers, Brownian motion in a field of force and the diffusion model of chemical reactions, Physica **7**, 284 (1940).

[2] P. Hänggi, P. Talkner, and M. Borkovec, Reaction-rate theory: fifty years after kramers, Reviews of modern physics **62**, 251 (1990).

[3] J. Miller, R. C. Petersen, E. Metter, C. Millikan, and T. Yanagihara, Transient global amnesia: clinical characteristics and prognosis, Neurology **37**, 733 (1987).
